# Analysis of gene network bifurcation during optic cup morphogenesis in zebrafish

**DOI:** 10.1101/2020.05.28.121038

**Authors:** Lorena Buono, Silvia Naranjo, Tania Moreno-Marmol, Berta de la Cerda, Rocío Polvillo, Francisco-Javier Díaz-Corrales, Ozren Bogdanovic, Paola Bovolenta, Juan-Ramón Martínez-Morales

## Abstract

Sight depends on the tight cooperation between photoreceptors and pigmented cells. Both derive from common progenitors in which a single gene regulatory network (GRN) bifurcates into the neural retina (NR) and retinal-pigmented epithelium (RPE) programs. Although genetic studies have identified upstream nodes controlling these networks, their regulatory logic remains poorly investigated. Here, we characterize transcriptome dynamics (RNA-seq) and chromatin accessibility (ATAC-seq) in segregating NR/RPE populations in zebrafish. Analysis of active cis-regulatory modules and enriched transcription factor (TF) motives suggest extensive network redundancy and context-dependent TF activity. Downstream targets identification highlights an early recruitment of desmosomal genes in the flattening RPE, revealing Tead factors as upstream regulators. Investigation of GRNs dynamics uncovers an unexpected sequence of TF recruitment during RPE specification, which is conserved in humans. This systematic interrogation of the NR/RPE bifurcation should improve both genetic counselling for eye disorders and hiPSCs-to-RPE differentiation protocols for cell-replacement therapies in degenerative diseases.

## Introduction

Darwin in *The Origin of Species* already hinted to the possibility of a single photoreceptor escorted by a pigmented cell, as the ancestral structure to all animal eyes. Several authors have further elaborated this notion during the last decades (Gehring, 2014; Vopalensky and Kozmik, 2009; Arendt and Wittbrodt, 2001). An intimate association between photoreceptors and pigmented ancillary cells is found in the eyes of all metazoan groups (Land and Nilsson, 2002), even in cnidarians (Kozmik, et al., 2008). Thus, the origin of this cellular tandem can be traced back to a common ancestor dating to at least 650 mya. Such remarkable evolutionary conservation can be explained by the fact that vision depends on the close physical and physiological interaction between the two cell types (Strauss, 2005). Photoreceptors, with a very active metabolism and a constant exposure to light, are particularly prone to degeneration due to DNA damage and oxidative stress. Pigmented cells protect photoreceptors by maintaining their homeostasis through growth factors secretion, visual pigments recycling and outer-segments phagocytosis (Letelier, et al., 2017).

Fate map experiments in vertebrates have shown that the precursors of the neural retina (NR) and retinal pigmented epithelium (RPE) derive from the undifferentiated eye field (Li, et al., 2000). In the permanently growing retinas of teleost fish, stem cells located at the ciliary marginal zone are capable of producing both cell types (Tang, et al., 2017). Furthermore, in tetrapods retinal precursors retain certain potential to trans-differentiate from the pigmented to the neural retina identity (Pittack, et al., 1991; Coulombre and Coulombre, 1965), and *vice versa* (Rowan, et al., 2004).

In zebrafish, the RPE and NR presumptive domains start differentiating at the optic vesicle stage from the medial (ML) and lateral (LL) epithelial layers, respectively (Kwan, et al., 2012; Li, et al., 2000). The specification of both retinal domains occurs simultaneously to the folding of the vesicle into a bi-layered cup, and entails profound cell shape changes in each domain. Precursors at the LL elongate along their apico-basal axis, differentiate as NR, and constrict basally to direct the folding of the retinal neuroepithelium (Nicolas-Perez, et al., 2016; Ivanovitch, et al., 2013). In contrast, precursors at the ML either acquire a squamous epithelial shape and differentiate as RPE, or flow into the LL to contribute to the NR domain (Moreno-Marmol, et al., 2018; Sidhaye and Norden, 2017; Heermann, et al., 2015; Picker, et al., 2009). In other vertebrates, the specification of the NR/RPE domains and the morphogenesis of the optic cup follow a pattern similar to that of the zebrafish, though the relative weight of the different morphogenetic mechanisms may vary among species (Martinez-Morales, et al., 2017).

During the last decades numerous studies have investigated how the RPE and NR domains get genetically specified (Fuhrmann, 2010). Eye identity is established in the anterior neural plate by the early activation of a gene regulatory network (GRN) pivoting on a few key regulators collectively known as eye field transcription factors (EFTF): including Lhx2, Otx2, Pax6, Rx, or Six3 (Zuber, et al., 2003). Upon the influence of inductive signals derived from the lens epithelium (FGFs) or the extraocular mesenchyme (Wnts and BMPs), the eye field network branches into the mutually exclusive developmental programs of the NR and RPE (Cardozo, et al., 2020; Fuhrmann, 2010). Classical genetic experiments performed mainly in mice have identified key nodes of the NR and RPE specification networks. The TF-encoding gene *Vsx2* (previously known as *Chx10*) was identified as the earliest determination gene differentially expressed in NR but not in RPE precursors (Liu, et al., 1994). *Vxs2* plays an essential role in specifying the NR domain by restraining the RPE identity through direct repression of the TF Mitf (Zou and Levine, 2012; Bharti, et al., 2008; Horsford, et al., 2005; Rowan, et al., 2004). Other homeobox regulators, many inherited from the eye field specification network, contribute to the network of NR specifiers. This list includes *Lhx2, Sox2, Rx, Six3* and *Six6* genes, which are required either for NR maintenance or for suppressing RPE identity (Wang, et al., 2016; Roy, et al., 2013). The establishment of the RPE network depends instead on the cooperative activity between Mitf and Otx factors (Lane and Lister, 2012; Martinez-Morales, et al., 2003). Classical experiments in mice showed that inactivation of either *Mitf* or *Otx* genes results in an RPE differentiation failure, with the alternative acquisition of NR molecular and morphological features (Martinez-Morales, et al., 2001; Nguyen and Arnheiter, 2000; Mochii, et al., 1998). More recently, mutation of the main effectors of the Hippo pathway, *Yap* and *Taz*, showed the essential role of these co-regulators in the differentiation of the RPE lineage (Kim, et al., 2016; Miesfeld, et al., 2015).

Despite the identification of these key regulators, the architecture of the NR and RPE specification subnetworks is far from being well understood (Martinez-Morales, 2016). Systematic attempts to reconstruct retinal GRNs using next-generation sequencing (NGS) methods have focused mainly in the differentiation of neuronal types at later stages of development or in epigenetic changes linked to retinal degeneration (Buono and Martinez-Morales, 2020; Wang, et al., 2018; Aldiri, et al., 2017). Very recently, scRNA-seq has been proved a useful tool to characterize cell heterogeneity and infer differentiation trajectories in human tissues and retinal organoids (Collin, et al., 2019; Hu, et al., 2019). However, a comprehensive analysis of the NR and RPE bifurcating networks would require a deeper sequencing coverage approach, while dealing with the limited cell population size at optic cup stages.

In this study, we dealt with previous limitations by applying a combination of RNA-seq and ATAC-seq on sorted NR and RPE populations. Using the zebrafish retina as a model system, we followed transcriptome dynamics and chromatin accessibility changes in both cell populations, as they depart from a common pool of progenitors. The cross-correlation of RNA-seq and ATAC-seq data allowed us to identify activating and repressing cis-regulatory elements (CREs), discover enriched transcription factor (TF) binding motifs, and unveil relevant downstream targets for the main specifiers. Our analyses confirmed previously known TFs as central nodes of the eye GRNs, and more importantly, provide information on their recruitment sequence both in zebrafish and human cells.

## Results

### 1 Analysis of specification networks in isolated NR and RPE precursors

To examine the bifurcation of the regulatory networks specifying the NR and RPE domains in zebrafish, we focused on the developmental window comprised between stages 16 hpf and 23 hpf (Figure 1 A). Within this period, the optic vesicle transits from a flattened disk in which the lateral and medial layers are still similar in terms of cell shape and volume, to an optic cup stage in which the NR and RPE cells are morphologically differentiated: bottle-shaped for retinal neuroblasts and flat for RPE precursors (Li, et al., 2000). It is important to state that at the end of this window (23 hpf), neurogenesis have not started yet in the retina, neither the RPE display pigmentation (Masai, et al., 2000). We used the transgenic lines *E1_bHLHe40:GFP* and *vsx2.2:GFP* to isolate the NR and RPE populations at 18 and 23 hpf by FACS (see methods; Figure 1B). Both *vsx2* and *bhlhe40* are among the earliest markers reported in zebrafish for the neural retina and RPE domains respectively (Cechmanek and McFarlane, 2017; Barabino, et al., 1997). To isolate early precursors at stage 16 hpf we took advantage of the fact that the *vsx2.2:GFP* transgene is transiently expressed in all progenitors and gets restricted to the NR as progenitors of the medial layer start expressing *bHLHe40* (Cechmanek and McFarlane, 2017; Nicolas-Perez, et al., 2016). Populations isolated by flow cytometry were analysed by RNA-seq, at all stages, and by ATAC-seq at 23 hpf (Figure 1A).

**Figure 1:**
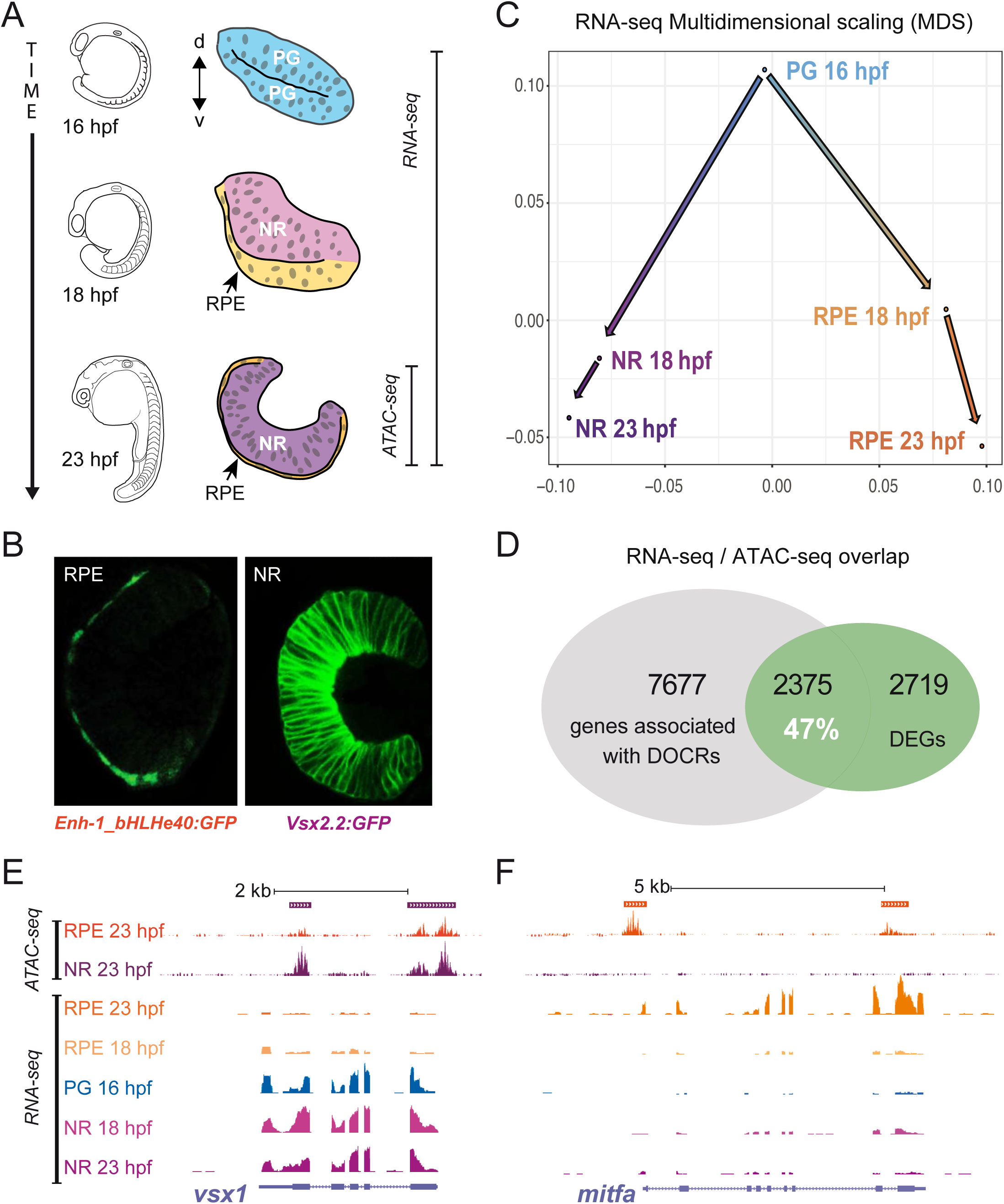
Experimental setup and raw data. (A) Schematic representation of optic vesicle morphogenesis from undiffe-rentiated retinal progenitors cells (PG, blue), at 16 hpf, to segregation and differentiation of the neural retina (NR, purple) and pigmented epithelium (RPE, orange) domains at 18 and 23 hpf. NR and RPE populations isolated by flow cytometry were analysed by RNA-seq and ATAC-seq at the stages indicated. (B) Stage 23 hpf confocal sections from the zebrafish transgenic lines used to mark and isolate the RPE (E1_bHLHe40:GFP) and NR (vsx2.2:GFP) populations. (C) Multidi-mensional scaling analysis of the RNA-seq data showing a progressive transcriptomic divergence between the NR and RPE domains. (D) Percentage of overlapping between differentially expressed genes (DEGs) and genes associated with differentially opened chromatin regions (DOCRs). (E-F) Overview of ATAC-seq and RNA-seq tracks (UCSC browser) for representative NR (vsx1; E) and RPE (mitfa; F) specific genes. Solid bars on the top indicate DOCRs. If purple, the DOCR is more accessible in the NR. On the contrary, if orange, the DOCR is more accessible in the RPE. In the depicted case all the DOCRs are accompanied by an increased transcription of the associated gene in the corresponding tissue.

The transcriptomic analysis of the different cell populations highlighted thousands of differentially expressed genes (DEGs) throughout development and between domains (Figure S1A,B; Dataset S1). A multidimensional analysis of these changes revealed that an extensive divergence between the NR and RPE transcriptomes already occurs within the first two hours (16 to 18 hpf) of optic vesicle folding (Figure 1C). Later on, between 18 hpf and 23 hpf, the transcriptome divergence between the NR and RPE progresses, although is more modest within each domain (Figures 1C and S1B).

### 2 Chromatin accessibility in NR vs RPE precursors identifies cis-regulatory modules

To gain further insight into the architecture of the optic cup GRNs, we sought to identify differentially open chromatin regions in the morphologically divergent NR and RPE domains. To this end, we performed ATAC-seq experiments from isolated NR and RPE precursors at 23 hpf. A total of 238369 open chromatin regions (OCRs) were detected using this approach. After statistical analysis, a fraction of the peaks was identified as differentially open chromatin regions (DOCRs), corresponding to putative cis-regulatory elements (CRE) that are more active in the NR than in the RPE or vice versa. Approximately 12.6% of all peaks (30172 peaks) were differentially open with an adjusted p-value < 0.05. This proportion was reduced to 4.8% when the adjusted p-value was lowered to <0.001 (Figure S2A,B; Dataset S2). Regardless the adjusted p-value, we observed a larger number of DOCRs in the RPE than in the NR (18909 vs 11263; adjusted p-value <0.05), and this was also accompanied by a higher average fold change in the RPE than in the NR associated DOCRs (Figure S2B,C). Analysis of the distribution of open chromatin regions in the genome showed no evident differences between NR and RPE for the whole set of OCRs. However, there was a difference in average localization between the entire OCRs and the subsets of DOCRs in the genome. When compared to the distribution of all the OCRs a noticeable depletion of DOCRs located near the promoter regions was evident in both the NR and RPE domains. The proportion of NR peaks near the promoter was 11.91% for OCRs vs 4.6% for DOCRs; whereas for the RPE peaks the proportion was 10.03% for OCRs vs 3.09% for DOCRs (Figure S2D), showing that domain specific cis-regulatory modules tend to occupy more distal positions in the genome. Gene ontology analysis of terms from the category Biological Process enriched in the list of genes associated with DOCRs yield results consistent with the analysed tissue. Genes associated with NR DOCRs were enriched in terms related to nervous system development, neuron differentiation, and eye morphogenesis, whereas genes associated with RPE DOCRs were enriched in terms such as melanocyte differentiation and epithelial differentiation (Figure S3).

Previous reports have shown the importance of correct DNA methylome patterning during human iPSC reprograming towards RPE (Araki, et al., 2019), and have proved a requirement for active demethylation during eye formation in vertebrates (Xu, et al., 2012). Thus, to further characterize the NR and RPE specific DOCRs, we next interrogated their DNA methylome profiles. We utilised base-resolution DNA methylome (WGBS) and hydroxymethylome (TAB-seq) data of zebrafish embryogenesis and adult tissues (Bogdanovic, et al., 2016), to obtain insight into the temporal dynamics of DNA methylation (5mC) over these open chromatin regions. Supervised clustering (k=2) of 5mC patterns separates both datasets into two well-defined clusters (Figure S4A,B). The first cluster consists of distal regulatory regions that initiate active demethylation at ∼24hpf, as demonstrated by strong 5-hydroxymethylcytosine (5hmC) enrichment and the progressive loss of 5mC starting at 24hpf, and then are most visible in the adult brain. Importantly, this cluster is highly enriched in H3K27ac, an active enhancer mark, and depleted of H3K4me3, which indicates their identity as bona fide enhancers (Bogdanovic, et al., 2012). The second cluster of both NR and RPE datasets is hypomethylated, depleted of hmC, and enriched in the promoter mark H3K4me3, in line with a CpG island promoters character. To test whether active, tet-protein dependent DNA demethylation is required for the establishment of open chromatin structure at NR and RPE ATAC-seq peaks, we interrogated chromatin accessibility in wild type and triple tet (tet1/2/3) morphants in whole 24hpf embryos (Bogdanovic, et al., 2016). Data from two independent MO knockdown experiments revealed that their chromatin accessibility signature is significantly (Mann-Whitney-Wilcoxon Test, P < 2.2e-16) reduced for NR ATAC-seq peaks in tet morphants when compared to wild type embryos, even when whole embryos were assessed (Figure S4C). This decrease was not visible in RPE peaks, likely because RPE enhancers are active in a much smaller cell population and their chromatin mark is masked by the en masse approach (Figure S4D).

### 3 Cross-correlation of RNA-seq and ATAC-seq data reveals activating and repressing CREs

Integration of ATAC-seq and RNA-seq data has been used as a powerful tool to explore the architecture of developmental GRNs (Buono and Martinez-Morales, 2020; Lowe, et al., 2019). By intersecting RNA-seq and ATAC-seq datasets, we observed a substantial overlap between DEGs and genes associated with DOCRs at 23 hpf. Thus, 47% of all DEGs are associated with at least one DOCR (Figure 1D). Specific examples of ATAC-seq and RNA-seq tracks are shown for the *vsx1* and *mitfa* loci, NR and RPE respective markers (Figure 1E,F). The cross-analysis of ATAC-seq and RNA-seq data allowed us to classify DOCRs (putative CREs) in two groups: those correlating with up regulated genes, here termed “activating CREs” and those correlating with silenced genes, here termed “repressing CREs” (Figure 2A,B). Illustrative examples of activating and repressing peaks are provided for the *six3a* and *otx2* loci (Figure S5A,B). When their average distance to the closest Transcription Start Site (TSS) was examined, we observed a significant trend: both for NR and RPE peaks, activating CREs tend to occupy positions closer to the TSS than the repressing regions (Figure S5C). A possible explanation for this difference is the positive bias introduced by proximal promoters, which are typically enriched in binding sites for activators (Maston, et al., 2006). To gain insight into the regulatory logic of the NR and RPE domains, we calculated the number of associated activating and repressing CREs for both the whole DEGs and the subset of differentially expressed TFs. Notably, as expected from their regulatory complexity, the average number of CREs per gene (either activating or repressing) was consistently higher for TFs than for the whole collection of DEGs (Figure 2C). More importantly, in the neural retina TFs associated with activating CREs outnumbered TFs associated with repressing regions (182 vs. 69 respectively), whereas the opposite was observed in the RPE. Indeed, in this tissue we detected a robust repressive cis-regulatory logic, with 173 TFs associated with at least one repressing CREs and only 110 associated with activating ones (Figure 2B,C). This opposite trend, better appreciated in histogram graphs, suggests that the NR specification program is sustained mainly by the activation of transcriptional regulators, whereas RPE determination seems to require primarily the repression of the NR-specific-TFs (Figure 2D,E).

**Figure 2:**
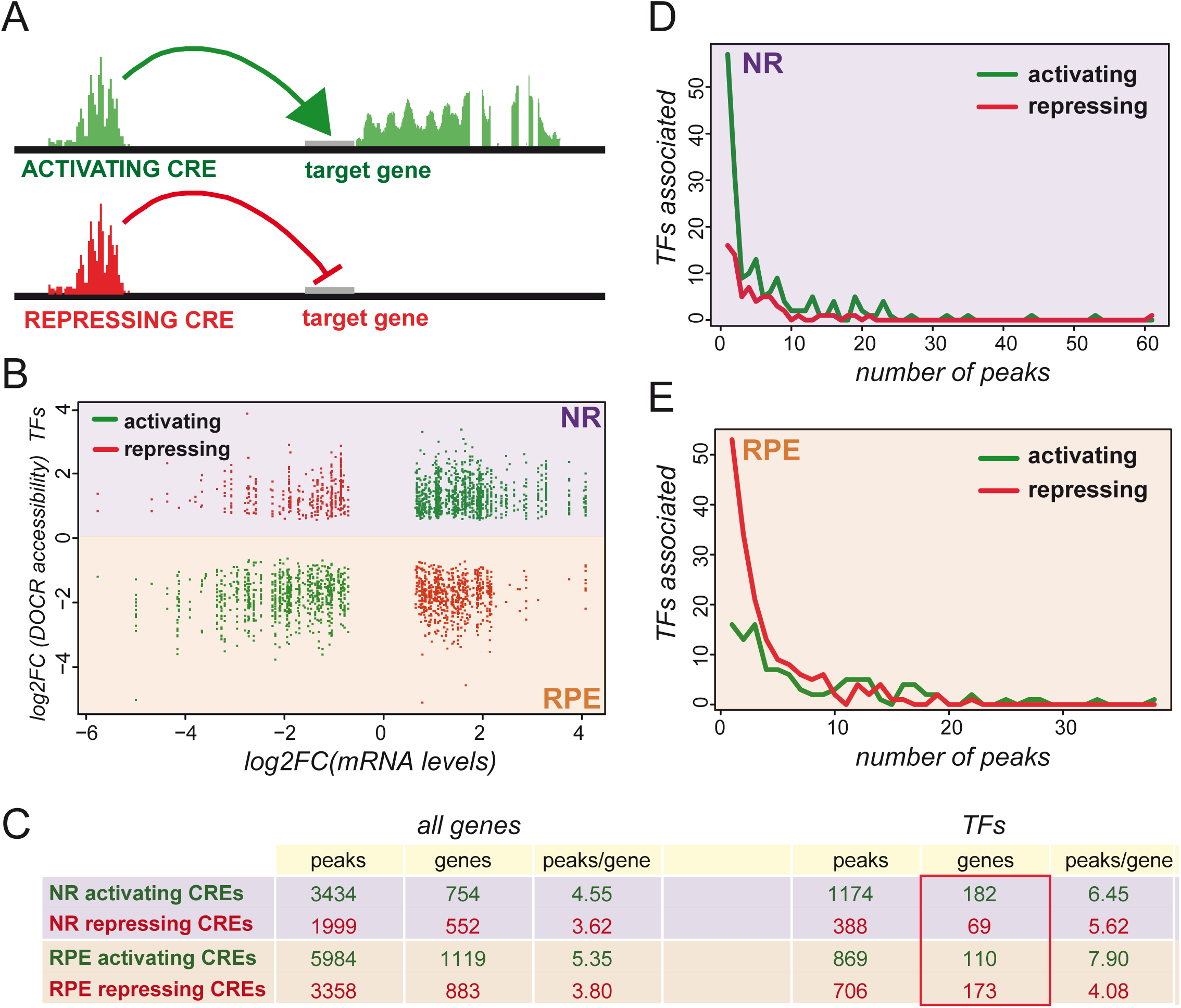
CRE configuration in the NR and RPE domains. (A) Schematic representation of the functional relationship between DOCRs, either activating (green) or repressing (red), and their associated DEGs. (B) The graph illustrates the correlation between the levels of differentially expressed TFs (log2FC) and the accessibility of their associated DOCRs (log2). (C) The table summarizes the number of activating or repressing CREs associated with either to all the DEGs or only with the differentially expressed TFs. (D-E) Histograms correlating the number of TFs associated with activating or repressing CREs to the number of peaks per gene, for both the NR (D) and RPE (E) domains.

### 4 Distinctive sets of TFs and cytoskeletal components are progressively recruited during NR and RPE specification

To efficiently analyse expression dynamics, as the NR and RPE networks bifurcate, we used a gene clustering approach. We applied both hierarchical and partitioning soft clustering for the classification of genes encoding for TFs or cytoskeletal components. We focused on these two gene categories to examine not only transcriptional lineage specifiers, but also terminal effectors operating in the divergent cell shape changes observed between the NR and RPE domains. Using partitioning soft clustering we established 25 groups of gene expression variation from progenitors towards NR or RPE, for both TFs and cytoskeleton components (Figure S6A,B; Datasets S3 and S4). This approach identified several expression clusters, which are distinctive for each domain and developmental stage: e.g. clusters 1, 13, and 23 for 16hpf progenitors in the TFs category (Figure S6A); or 6, 11, and 21 for 23hpf RPE precursors in the cytoskeletal components’ analysis (Figure S6B). Using a hierarchical clustering approach, we could aggregate the small sub-groups into 6 main clusters for TFs and 8 for cytoskeletal components, all of them linked mainly to a specific domain and/or developmental stage (Figures 3 and S7; Datasets S5 and S6). To define precise time windows of expression and to infer regulators order of action, we focused our attention on the identity of TFs belonging to the different large clusters. Significant TFs in cluster #5, corresponding to 16hpf progenitors, include *rx3*, an early eye specifier with a known role in optic vesicle evagination (Rembold, et al., 2006; Loosli, et al., 2001; Mathers, et al., 1997); and *her* factors required to maintain the progenitor neural state (Chapouton, et al., 2011) (Figure 3B). As expected, Cluster #1, corresponding to NR 23 hpf precursors, contains many of the acknowledged retinal specifiers, such as *rx1, rx2, sox2, six3a, six3b, six6b, vsx1, vsx2, hmx1, hmx4* and *lhx2b* (Fuhrmann, 2010), which already increased their expression at 18 hpf (Figure 3B). Surprisingly, known RPE specification genes such as *otx2* and *mitfa* (Martinez-Morales, et al., 2004), included in cluster #4, do not increase their levels significantly in RPE precursors at 18hpf, peaking only later at 23 hpf (Figure 3B). In contrast, TFs included in cluster #3, such as *tead3b, tfap2a* and *tfap2c* started to rise, when not peaking, in the RPE at 18 hpf. Notably, some of these TFs, including *tcf12, smad6b*, and especially *vgll2b* not only peak at 18 hpf, but also rapidly decrease at 23 hpf (Figure 3B). This observation suggests the existence of two waves of TFs regulating the identity of the RPE domain.

**Figure 3:**
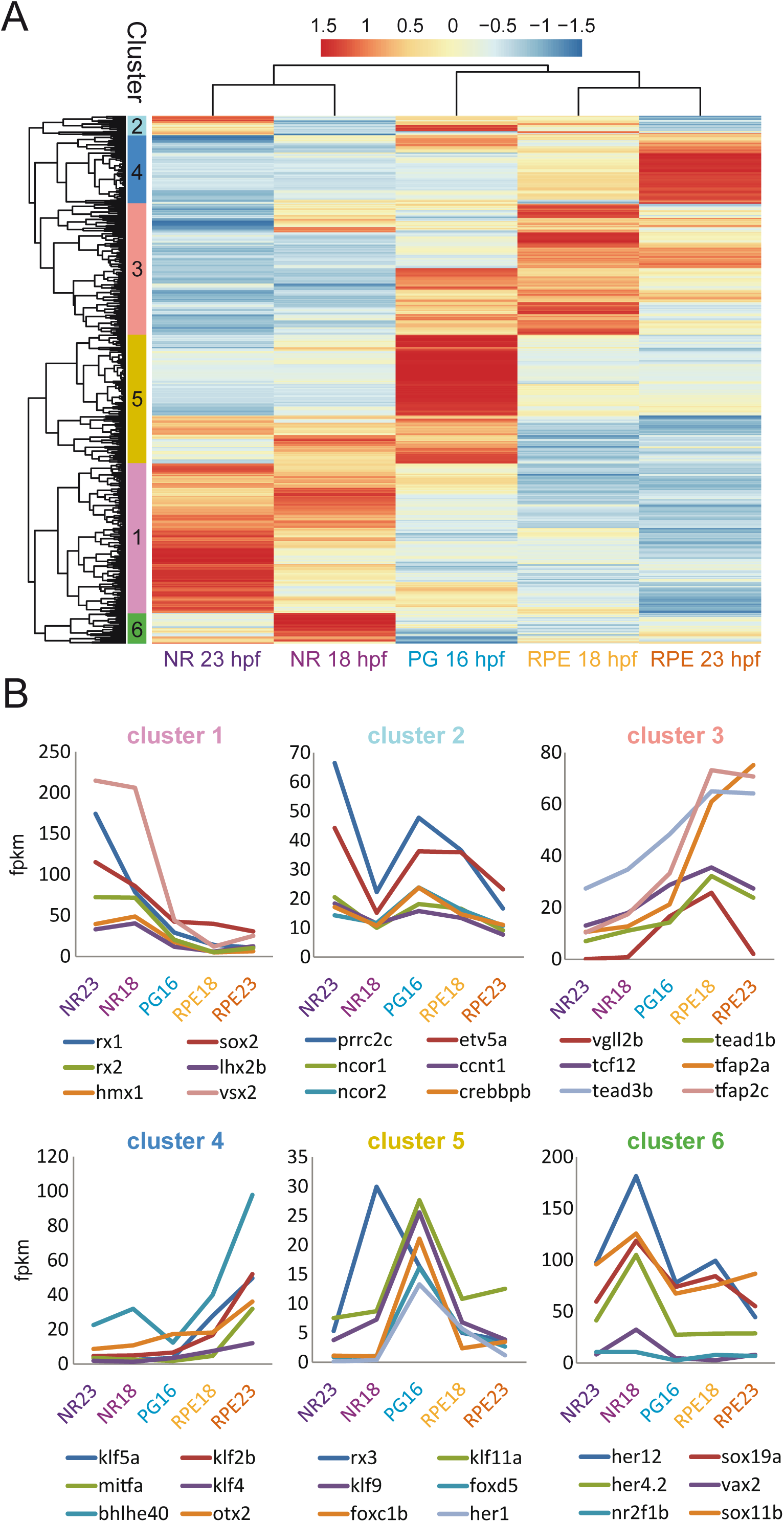
Hierarchical clustering of TFs gene expression variations during optic cup development. (A) Hierarchical clustering output shows TFs expression trends in the distinct domains and stages. Gene expression values, normalized by row, are indicated with a red to blue graded colour. Note that most clusters comprise a particular domain and developmental stage. (B) Relative transcript level changes for significant TFs (indicated by dotted lines) within each cluster.

Next, we focused on the content of the hierarchical clusters associated to cytoskeletal components (Figure S7A). While the analysis of the cytoskeletal genes upregulated in PG returned no significant enrichment for terms annotated in the Cellular Components, similar analysis for cytoskeletal genes in the NR clusters yielded significantly enriched terms associated to microtubules and centrosomes (Figure S7B). This observation is in line with the elongation of the apico-basal axis and the polarization of the microtubules reported for NR precursors in zebrafish (Ivanovitch, et al., 2013). In contrast, the only enriched term in the RPE cytoskeletal clusters was “intermediate filaments” (Figure S7B), consistent with the RPE acquiring a squamous epithelial character.

### 5 Motif enrichment analysis suggests redundancy and cooperativity in the NR and RPE networks

To further explore the regulatory logic of the NR and RPE gene networks, we investigated overrepresented motifs within the DOCRs associated with each domain. Motif enrichment analysis of the differentially open regions in the NR identified a highly significant overrepresentation of the homeobox and sox TF binding motifs (Figure 4A; Dataset S7). The core homeobox binding motif (5’-TAATT-3’) is shared by TFs from the homeodomain K50 PRD-class; such as *vsx1, vsx2, rx1, rx2*, and *rx3*; LIM-class, such as *lhx2b*; and NKL class; such as *hmx1* and *hmx4* (Figure 4A). All of them are well-known retinal specifiers contained in group #1 of our hierarchical clustering analysis (NR 23 hpf). As they are co-expressed in the retina, it is likely that they cooperate to target a partially overlapping set of cis-regulatory modules. To explore this possibility, we retrieved the individual position weight matrixes (PMW) associated to both homeobox and sox TFs from available databases and used this information to explore their functional synergy in the retinal DOCRs (see Method sections). Two different approaches were used: (i) calculating the co-occurrence rate of binding sites for two different TFs in the same CRE (co-occupancy); and (ii) estimating the percentage of binding sites for TFs located in different CREs but still associated with the same gene (co-regulation) (Figure 4B,C). These analyses predicted an extremely high degree of inter-connectivity within the NR network. Thus, the average co-occurrence rate of two different theoretical binding sites in the same peak (co-occupancy) was 25.6%. This combinatorial activity becomes more pronounced when examining the binding sites in different CREs associated to the same gene (co-regulation), which was in average 46.6%. Indeed, when individual TF binding sites were investigated, the number of co-occupied peaks or co-regulated genes was in general much larger than those instances in which the binding site was found isolated (Figure 4D,E). Even when these are predicted binding sites, our analyses suggest a strong cooperative activity of homeobox and sox factors in driving the NR developmental program.

**Figure 4:**
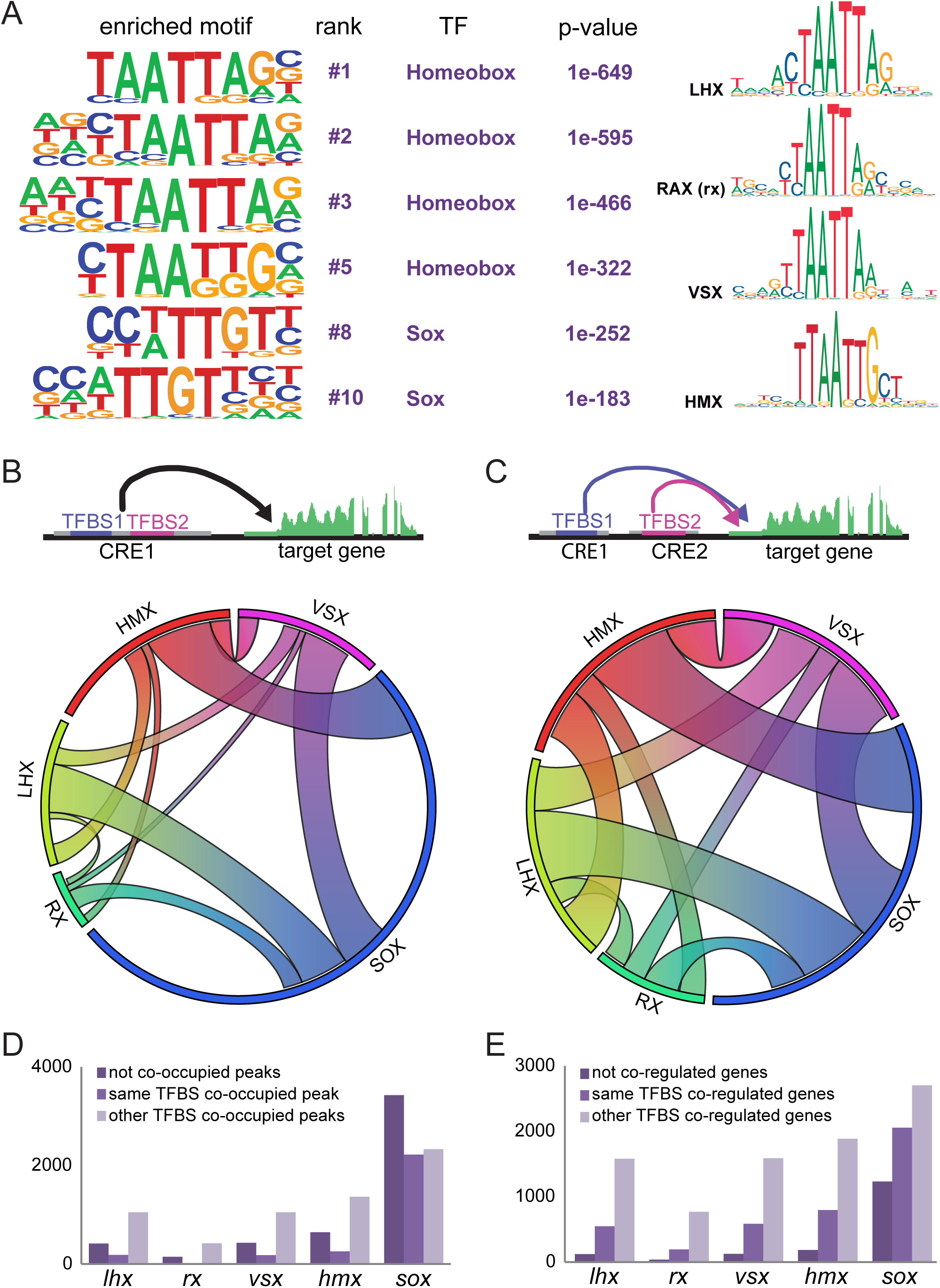
NR motif enrichment analysis. (A) Representative TF binding motifs enriched in NR DOCRs as identified by HOMER. The binding motif similarity among neural retina TFs of the homeobox family is indicated (http://jaspar.genereg.net). (B) Circoplot illustrating the co-occupancy rate of TFBS in the same DOCRs for the main TFs identified in the motif enrichment analysis (C) Circoplot illustrating the degree of co-regulation between TFs regulating the same gene through different DOCRs. (D) Number of CREs containing the main TFs identified in the motif enrichment analysis, classified according to their co-occupancy (E) Number of genes associated to CREs containing the main TFs identified in the motif enrichment analysis, classified according to their co-regulation.

In the RPE, a motif enrichment analysis of the specific DOCRs also revealed a number of binding motifs that correspond to the TFs identified as up-regulated in the RNA-seq analysis (Figure 5A; Dataset S8). We found a very significant enrichment for tfap2a and tfap2c binding motifs, in agreement with the early expression of these factors in the RPE. This considerable overrepresentation points to a prominent position of these factors within the regulatory hierarchy, which is in line with their role in the specification of the pigmented tissue (Bassett, et al., 2010). In addition, a significant enrichment for bHLH, tead and otx2 binding motifs was also observed. Importantly, the core bHLH motif (5’-CACGTG-3’) is shared by members of the family expressed in RPE, such as *mitfa, bHLHE40, bHLHE41*, and *tfec* (Figure 5A). As already investigated for the NR TFs, we examined the potential cooperation among these RPE specifiers (Figure 5B-E). Interestingly, whereas the average rate of co-regulated genes in the RPE did not differ much from that found in NR (48% vs 46.6%; Figure 5E), the rate of co-occupied peaks dropped from 25.6% in the NR to 11.7% in the RPE (Figure 5D). This observation suggests that the RPE network is less dependent on TFs cooperativity within the same CRE than the NR network. In such regulatory scenario, each TF may trigger transcriptional sub-programs within a broader developmental network. To explore this possibility, we assessed GO enrichment for the genes associated with RPE DOCRs containing the different TFBS and then we grouped the results using hierarchical clustering (Figure S8). The outcome of this clustering approach supported a branched regulatory scenario. Thus, whereas some TFs (i.e. *tcf12, tfap2c, otx2*) appeared associated to many GO terms, others were more specific: such is the case of *mitfa* that was associated only to pigment cell differentiation (Figure S8).

**Figure 5:**
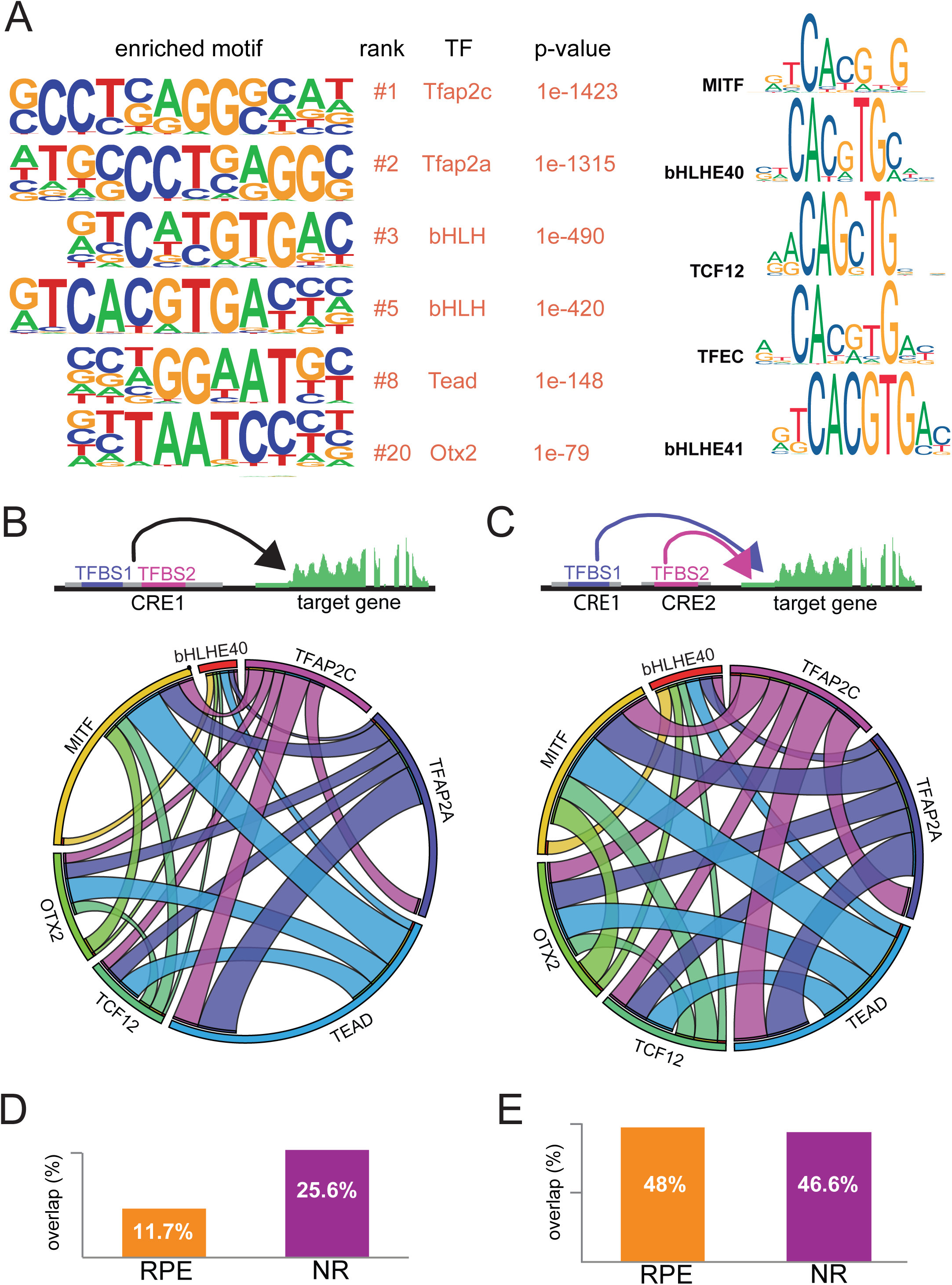
RPE motif enrichment analysis. (A) Representative TF binding motifs enriched in RPE DOCRs as identified by HOMER. Analysis of the binding motif similarity among TFs of the bHLH family (http://jaspar.genereg.net). (B) Circoplot illustrating the co-occupancy rate of TFBS in the same DOCRs for the main TFs identified in the motif enrichment analysis (C) Circoplot illustrating the degree of co-regulation between TFs regulating the same gene through different DOCRs. (D) Average percentage of co-occupancy in the same DOCR for two different TF in the RPE and NR. Average percentage of two different TF regulating the same gene through different DOCRs in the RPE and NR.

Finally, to investigate the regulatory logic of activating and repressing CREs in each domain, we scanned these regions for enriched motifs separately. Strikingly, those enriched motifs ranking higher for the NR activating CREs, such as the homeobox and sox TF binding sites, were also ranking higher for the repressing regions in this tissue (Figure S9; Dataset S9). A very similar scenario was observed for the RPE peaks, which regardless being activating or repressing regions showed tfap2c, tfap2a, bHLH, and otx2 consensus binding sites as top enriched motifs. When GO terms associated with genes linked to these regions were examined, we observed terms linked to eye morphogenesis enriched for both NR activating and RPE repressing regions (Figure S9). This finding reveals context-dependent TFs activity, and points to a common set of genes activated in the NR and repressed in the RPE by antagonistic GRNs. In fact, a detailed analysis of the gene lists associated to NR activating and RPE repressing regions showed that many of the retinal specifiers themselves (including *hmx4, lhx9, mab21l1, nr2f2, pax6a, pax6b, rx2, six3a*, or *sox2*) are under the antagonistic regulation of the NR and RPE GRNs (Dataset S10). In addition, the analysis of GO terms enrichment also suggested domain-specific functions for the NR and RPE GRNs. Thus, NR repressing CREs are associated to genes involved in mesoderm formation, whereas RPE activating regions are link with epidermal differentiation genes.

### 6 Desmosomal components are activated during RPE specification

The integration of our ATAC-seq and RNA-seq data allows formulating and testing different hypotheses related to genetic programs controlling the specification of retinal tissues. As an example, we followed up the observation that genes encoding for intermediate filaments are enriched in the RPE domain (Figure S7B). Keratin looping into desmosomal plaques plays a fundamental role in maintaining tissue architecture under mechanical load (Hatzfeld, et al., 2017) (Figure 6A). We thus investigated the transcriptional profile of other desmosomal components as the RPE and NR networks diverge. A detailed analysis revealed that not only keratin genes, but also many desmosomal genes such as *dspa, evpla, pleca*, or *ppl*, are among the most upregulated genes (i.e. highest fold change) in committed RPE cells (Figure 6B). Many of these genes increased their expression already at 18 hpf, when RPE cells start to differentiate morphologically, but hours before the tissue acquires pigmentation (Figure 6B). To gain insight into the regulation of these cytoskeletal components, we performed a motif enrichment analysis of the subset of DOCRs associated with keratins and other desmosomal genes upregulated in the RPE (Dataset S11). This analysis revealed Tead motifs as top ranked overrepresented binding sites in the CREs regulating desmosomal genes (Figure 6C). When compared to that of the entire set of DOCRs, the average number of motifs per peak was 4.5 fold higher for those CREs specifically associated with components of the desmosome machinery (1.35 vs 0.3). This observation suggests a key role for Tead family proteins in the regulation of the intermediate filament cytoskeleton. To test this hypothesis, we took advantage of available zebrafish double mutants for the main Tead coactivators *yap* and *taz*, which have been reported to display RPE differentiation defects (Miesfeld, et al., 2015). In agreement with our observations, the expression of keratins (*krt4* and *krt8*) as well as pigmentation genes (*tyr* and *tyrp1b*) were significantly reduced in *yap -/- taz -/-* double mutant head tissue at 18 hpf as determine by RT-qPCR (Figure 6D). As a control, the expression of the NR markers *vsx2* and *six3a* was unaffected in the mutant tissue. These results further support a role for Tead factors in the transcriptional regulation of desmosomal genes at the RPE.

**Figure 6:**
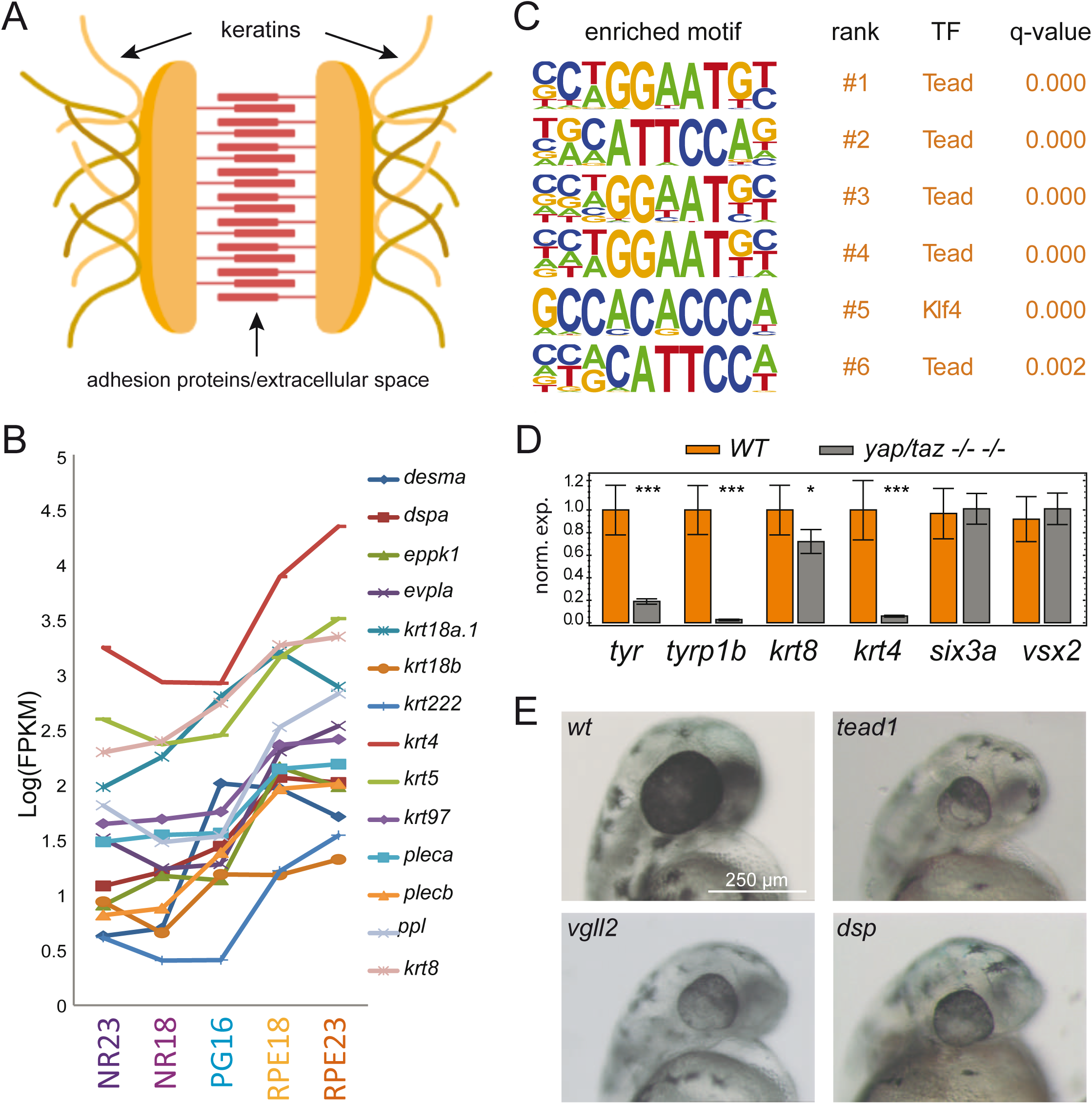
Regulation of desmosomal components during RPE specification. (A) Schematic representation of a desmosome junction. (B) Intermediate filament and desmosomal genes expression variations during optic cup development. Expression values are reported as log(FPKM) (C) Motif enrichment analysis of the DOCRs associated with genes encoding intermediate filament or desmosome components. (D) mRNA levels of keratin genes as well as RPE and NR markers as determined by RT-qPCR in wild type and yap -/- taz -/- double mutant zebrafish samples (dissected heads) at 18 hpf. Significant differences are indicated (n=3; T-test; ***= p<0.001). (E) Representative stereo microscope images of zebrafish embryo heads at 48 hpf. Magnification Bar = 0.5 mm): wild type and embryos injected with Cas9 (300 ng/μl) together with the following sgRNAs (80 ng/μl) combinations: vgll2a and vgll2b (vgll2); tead1a and tead1b (tead1) and dspa and dspb (dsp). Note the reduced eye size and RPE hypopigmentation in the crispants.

The high efficiency of the CRISPR/Cas9 technology allows performing F0 mutagenesis screens in zebrafish (Shankaran, et al., 2017). We took advantage of this technology to test functionally some of the genes identified in our analyses. A total of 21 genes, selected on the bases of their expression profile, CRE composition and/or dynamics, associated GO terms, and number of paralogues, were investigated in an F0 pilot screen (Figure S10). A substantial proportion of the tested genes (i.e. 12 out of 21 sgRNA combinations tested) displayed eye malformations, such as microphthalmia, eye fissure closure defects and/or hypopigmentation, thus confirming that many of the candidate genes play an essential role as components of the eye GRNs (Figure S10; Dataset S12). Interestingly, among the sgRNA combinations tested, the injection of those directed against the structural desmosomal components *dspa/b*, the Tead regulators *vgll2a/b*, and particularly the cocktail *tead1a/b* + *tead3a/b* resulted in a significant proportion (30.4%, 40.6%, and 80% respectively) of the F0 embryos displaying reduced eye size and severe hypopigmentation (Figures 6E and S10). These results further indicate that both desmosomal assembly and Tead activity are required during RPE differentiation.

### 7 Gene expression analysis during hiPSCs-to-RPE differentiation

A second important finding derived from our gene expression clustering analysis was the identification of two waves of transcriptional regulators during the specification of the RPE in zebrafish. Understanding this TF recruitment sequence in humans may have important basic and translational applications. We thus asked whether the same *consecutio temporum* identified in zebrafish was conserved in human iPSCs differentiating to RPE. During the first four weeks of differentiation in culture from hiPSCs (see methods), pluripotent stem cells progressively change their morphology to a cobblestone appearance, acquiring a light pigmentation at the end of the fourth week (Figure 7A). Using RT-qPCR, we tested mRNA levels for a total of 26 RPE genes including genes encoding for stemness markers (*NANOG* and *OCT4*); mature RPE markers (i.e. *CRALBP, RPE65* and *TYR*); known RPE specifiers activated in the second wave of gene expression (i.e. *BHLHE40, MITF, OTX2*, and *TFEC*); desmosomal components (i.e. *DSP, EVPL, KRT4, KRT5, KRT8*); and TFs and signalling molecules activated in the first wave of gene expression (NOTCH1, NOTCH2, NOTCH3, SMAD6, TCF12, TEAD1, TEAD2, TEAD3, TEAD4, TFAP2A, TFAP2C, and VGLL2) (Figure 7B; Dataset S13). This analysis showed that most of the human genes orthologous to the zebrafish early specifiers cluster together according to their expression profiles, and reach maximal expression within the first weeks of culture. This was the case for *SMAD6, TEAD1, TEAD3, TFAP2A, TFAP2C*, and particularly for *VGLL2 and TEAD2*, the expression of which peaked very transiently. Similarly, the human orthologs of most of the genes that in zebrafish were identified as activated in a second transcriptional wave (including *MITF, BHLHE40, TFEC, CRALBP, RPE65* and *TYR*) also clustered together reaching a maximum expression level in the fourth week of culture. These findings indicated that despite the very different time scales of the developmental programs in these far-related vertebrate species, hours in zebrafish and days in humans, their RPE regulatory networks share a common logic of two-waves in the TFs recruitment sequence.

**Figure 7:**
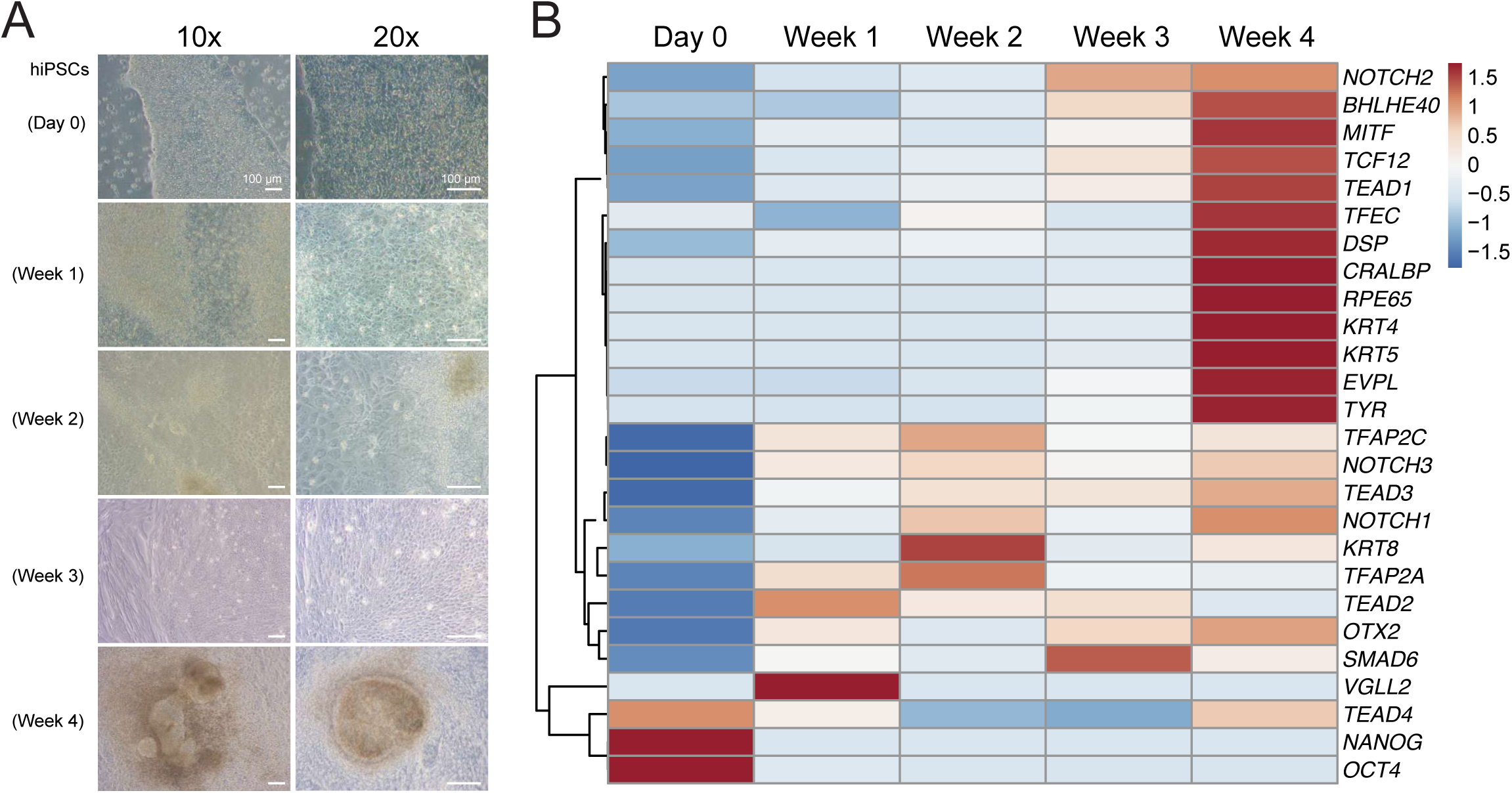
Gene expression during hiPSCs-to-RPE differentiation. (A) Bright field microscopy images (10-x and 20x) of hiPSCs before (Day 0) and during their differentiation towards RPE. Note the progressive acquisition of epithelial morphology and pigmentation. (B) Hierarchical clustered heatmap showing gene expression level variations, as determined by RT-qPCR during the differentiation towards RPE from hiPSCs. Note the conservation of the two clusters (early and late transcriptional waves) identified in zebrafish.

## Discussion

### A system level approach to explore GRNs architecture

Cell differentiation, as classically illustrated by Waddington’s epigenetic landscapes (Waddington, 1966), often entails sequential binary decisions that progressively restrict cell competence. At a genetic level this translates in the branching of “mother” gene regulatory networks into child sub-networks. Here we have used a double RNA-seq/ATAC-seq approach to zoom into the bifurcation of the NR/RPE developmental programs, which will give rise to specialized neurons and pigmented squamous cells respectively. The transient nature of this differentiation process, as well as the limiting size of the cell populations involved (Kwan, et al., 2012), have hindered any systemic approach to investigate the architecture of the specification networks as they branch. The high sensitivity and deep sequencing coverage of the RNA-seq and ATAC-seq methods (Buenrostro, et al., 2013), combined with the isolation of NR and RPE precursors by FACS allowed us to overcome previous constrains. Our approach permitted not only the detection of transcriptomic variations and active cis-regulatory modules, but also helped to define hierarchical relationships among the core components of the network. In our analyses we have used a proximity method to associate genes and open chromatin regions. Previous studies have confirmed that this link to the adjacent gene is correct in 90% of the cases (Yoshida, et al., 2019). Therefore, although this estimate should be taken cautiously when individual cis-regulatory elements are examined, it is a valid approach when conclusions are driven from a genome-wide analysis. In this study we identified approximately 30000 chromatin regions differentially open (DOCRs) between the NR and RPE domains. The examination of their methylation status, as well as their overlap with H3K27ac and H3K4me3 marks indicates that most of these regions (≈95%) correspond to initially inactive enhancers that get progressively demethylated and activated at the phylotypic period, a behaviour reported for developmental genes in general (Bogdanovic, et al., 2016). A much smaller cluster (≈ 5%) matches to hypomethylated constitutively active promoter regions (Figures S2 and S4). It is therefore likely that most of the regions here identified as DOCRs correspond to cis-regulatory elements (CREs) active during the segregation of the NR/RPE networks.

### The NR as a default program

Our data, together with previous observations by others, are consistent with the hypothesis that the neural retina is the default state of the optic vesicle precursors. Morphologically, optic vesicle precursors either at the medial or the lateral layers share a similar neuroepithelial character: i.e. elongated cells polarized along the apico-basal axis arranged in a pseudostratified epithelium (Ivanovitch, et al., 2013; Kwan, et al., 2012). During optic cup morphogenesis, NR precursors retain this neuroepithelial morphology, whereas differentiating RPE cells undergo profound cell shape changes as they progressively flatten into a squamous epithelium (Moreno-Marmol, et al., 2018). These observations correlate in our analyses with a larger number of differentially upregulated genes and differentially opened chromatin regions at the RPE (Figure S2). We have shown that repressing CREs associated with TFs dominate in the RPE network, whereas activating elements are more abundant in the NR program (Figure 2). Furthermore, the motifs analysis of activating elements in the NR and repressing elements in the RPE indicates that they are antagonistically regulating a similar set of genes (Figure S9). It is therefore very likely that the acquisition of the RPE fate requires the global repression of the NR program. However, although the role of *Vsx2* in the suppression of RPE identity via direct *Mitf* repression has been well documented (Zou and Levine, 2012; Horsford, et al., 2005; Rowan, et al., 2004), the mechanisms underlining the repressive activity of the RPE network are much less understood (Fuhrmann, et al., 2000). Interestingly, *Pax6* seems to cooperate both with NR and RPE specifiers acting as a balancing factor between the two networks depending on the cellular context (Raviv, et al., 2014; Bharti, et al., 2012). The complexity of the transcriptional regulatory logic is often exemplified by its context dependence, which may happen at very different levels. As illustrated by a recent report in *Drosophila*, cis-regulatory modules can play a dual role as enhancers or silencers in different cellular environments (Gisselbrecht, et al., 2020). Our motif analysis also points to the same set of TFs acting as repressors or activators within the same tissue, suggesting their activity is conditioned by their chromatin environment. This is in line with previous work showing that TFs have diverse regulatory function depending on the enhancer context (Stampfel, et al., 2015).

### GRNs robustness

One of our main observations is that NR and RPE networks are remarkably robust. This is the case particularly in teleosts, in which whole genome duplication resulted in additional copies of eye specifiers. In zebrafish, several homeobox TFs expressed in the neural retina converge on the 5’-TAATT-3’ motif, including vsx1, vsx2, rx1, rx2, rx3, lhx2b, lhx9, hmx1 and hmx4. Our motif discovery predictions suggest these proteins, together with sox factors, may coregulate the same genes and cooperate within the same CREs, although these are theoretical predictions that may require validation by direct ChIP-seq studies. This approach though is challenging in our experimental setting due to the limited starting material and the reduced availability of ChIP grade antibodies in zebrafish. Yet, we believe that considered in bulk these are informative predictions, as they are restricted to differentially open chromatin regions and differentially expressed TFs. Supporting a co-regulation, the mutation of many retinal homeobox TFs, such as *rx2* in medaka (Reinhardt, et al., 2015) or *lhx2* in zebrafish (Seth, et al., 2006) does not compromise severely the identity of the tissue. In the RPE, redundancy pertains mainly the core bHLH motif (5’-CACGTG-3’), which is shared by mitfa, bHLHE40, bHLHE41, and tfec. In agreement with this, the simultaneous elimination of *mitfa* and *mitfb* has no apparent consequences on RPE specification or pigmentation in zebrafish (Lane and Lister, 2012). All these observations further point to a broad redundancy and cooperativity among the main nodes of the eye specification networks.

### Coupling of transcriptional and cell shape changes in the RPE

As expected, the cell shape changes associated to the acquisition of the NR and RPE identities have a clear reflection at the transcriptomic level. The differentiation of the NR entails the activation of cytoskeletal components associated to microtubules polarization, in agreement with the neuroepithelial character of this tissue (Ivanovitch, et al., 2013). In contrast, here we show that a significant number of keratins and other desmosomal genes get recruited in the presumptive RPE hours before this tissue acquires its distinctive pigmentation. Although keratins and plaque proteins have been reported as RPE markers in several vertebrate species (Owaribe, et al., 1988), their contribution to the morphogenetic program of this tissue remains unexplored. Given the role of desmosomal plaques in conferring resistance to mechanical load (Hatzfeld, et al., 2017), it is tempting to hypothesize that the activation of these cytoskeletal genes may be linked to RPE precursors adapting their cytoskeleton to increased tissue tension as they flatten. Motif enrichment analysis of CREs linked to desmosomal genes, together with the genetic evidence here provided by *yap -/-*; *taz -/-* double mutants and Tead crispants, strongly support a role for Tead TFs in the activation of keratin genes at the RPE. Indeed, several keratin (e.g. *krt5, krt8*, and *krt97*), and desmosomal genes (e.g. *evpla, pleca, plecb*, and *ppl*) have been identified as targets for Yap/Tead complexes by ChIP-seq studies in mammalian cells (Estaras, et al., 2017; Zanconato, et al., 2015; Lian, et al., 2010), as well as DamID-seq studies in zebrafish embryos (Vazquez-Marin, et al., 2019). Thus, the direct transcriptional regulation of keratins and plaque genes by Yap/Tead complexes seems a conserved theme across different tissues and vertebrate species.

### Two-waves of TFs recruitment during RPE specification

Arguably one of the most notable observations in this study is the relatively late peak of expression of genes such as *mitfa* or *otx2*, which were previously considered among the earliest RPE specifiers (Martinez-Morales, et al., 2004). Their delayed peak of expression occurs after the specification and flattening of RPE precursors have commenced, and rather coincides with the onset of pigmentation in the tissue at 23hpf. This agrees with previous studies indicating a cooperative role for Otx and Mitf in the direct activation of the melanogenic gene battery (Martinez-Morales, et al., 2003). A detailed expression study in zebrafish also reported two separated phases for RPE specification, the first of which is *mitf* independent (Cechmanek and McFarlane, 2017). Here we show that the early phase of specification entails the recruitment of a different set of transcriptional regulators, including *smad6b, tead1b, tead3b, tfap2a, tfap2c, tcf12*, and *vgll2b*. Some of these TFs, or their cofactors, act as upstream regulators of *Otx* and *Mitf*, such is the case for *Tcf* (Westenskow, et al., 2009) and Tead-cofactors *Yap* and *Taz* (Kim, et al., 2016; Miesfeld, et al., 2015). Others, such as *tfap2a*, play an important role in RPE specification, although their hierarchical role within the RPE GRN remains unclear (Bassett, et al., 2010). Finally, here we also describe a similar sequence of TF recruitment in human differentiating RPE cells. Thus, despite the different morphology of the cells (i.e. cuboidal in humans and squamous in zebrafish) and the large evolutionary distance (≈450 mya), the regulatory logic that specifies the RPE seems largely conserved across vertebrates.

### Conclusions

Our data shed light on the bifurcation of the NR and RPE programs not only in zebrafish but in other vertebrate species. The obtained results may be relevant to identify novel causative genes for eye hereditary diseases, as mutations in many nodes of the eye GRNs result in congenital eye malformations (Fitzpatrick and van Heyningen, 2005). Importantly, our findings uncover an unanticipated regulatory logic within the RPE specification network. This provides critical information to improve hiPSCs-to-RPE differentiation protocols, a key step in cell replacement strategies for retinal degenerative diseases.

## Methods

### Fish maintenance

The zebrafish (Danio rerio) AB/Tübingen (AB/TU) wild-type strains, the transgenic lines tg(vsx2.2:GFP-caax) (Nicolas-Perez, et al., 2016) and tg(E1_bHLHe40:GFP) (Moreno-Marmol et al., 2020) and the mutant strain yap +/- taz +/- (Miesfeld, et al., 2015) were maintained and breed under previously described experimental conditions (Westerfield, 2000). All animal experiments were carried out according to the guidelines of our Institutional Animal Ethics Committee.

### Cell cytometry

Zebrafish cells were dissociated and prepared for FACS as previously described (Manoli and Driever, 2012). PG at 16 hpf, NR at 18 and 23 hpf were isolated from dissected heads of the tg(*vsx2.2:caax-GFP*). RPE at 18 and 23 hpf were isolated from whole tg(E1_*bHLHE40*:GFP) embryos. A FACSAriaTM Fusion flow cytometer was used to recollect only the GFP+ cells. GFP+ cells were isolated directly in Trizol for RNA extraction, or in ATAC-seq tagmentation buffer for open chromatin detection.

### RNA extraction

Total RNA was extracted using 750 ul TRIzol LS (Invitrogen) following manufacturer’s protocol. Possible DNA contamination was eliminated treating the RNA samples with with TURBO DNAse-free (Ambion). Concentration of the RNA samples was evaluated by Qubit (Thermo Fisher), and then the samples were used for subsequent applications.

### qPCR

cDNA retrotranscription and qPCR were performed as described (Vazquez-Marin, et al., 2019). For primer sequences, see Tables S1 and S2. *HPRT1* and *GAPDH* were used as housekeeping genes for human samples, whereas *eef1a1l1* was used for zebrafish samples.

### RNA-seq analyses

RNA was extracted from sorted cells and then treated with DNAse as described above. rRNAs were eliminated from the samples with Ribo-Zero^®^ rRNA Removal Kit (Illumina) prior library preparation. Samples were sequenced in SEx125bps or PEx125bps reads with an Illumina Hiseq 2500. We obtained at least 35 M reads from the sequencing of each library. Three biological replicates were used for each analyzed condition. Reads were aligned to the danRer10 zebrafish genome assembly using Tophat v2.1.0. Transcript abundance was estimated with Cufflinks v2.2.1. Differential gene expression analysis was performed using Cuffdiff v2.2.1, setting a corrected p value < 0.05 as the cutoff for statistical significance of the differential expression. Multidimensional Scaling Analysis (MDS) was performed using the function MDSplot of the CummeRbund package in R 3.6.1. Soft clustering of time-series gene expression data was done for all the transcripts with a variance among the five conditions ≥ 3 using the R package Mfuzz with a m = 1.5 (Kumar and M, 2007). The TF transcript subset was extracted from the total list of genes using the tool “Classification System” of PANTHER (Thomas, et al., 2003) filtering for the protein class “transcription factors” (PC00218). Some TFs not present in the database for an annotation issues (i.e. mitfa, vsx1, vsx2, rx1, rx2, rx3, lhx2b, hmx1, hmx4, sox21a) were added manually. The cytoskeleton component subset was obtained retrieving all the genes belonging to GO term “cytoskeleton” (GO:0005856), including all the child and further descendant GO terms, with biomaRt. All the heatmaps representing transcriptomic variations were plotted with the R package *pheatmap* using exclusively the transcripts that resulted to be differentially expressed from the comparison between at least two of our experimental conditions TFs and cytoskeleton components were filtered using the same methodology used for Mfuzz clustering. Gene ontology analysis was performed with the online tool GOrilla (Eden, et al., 2009) or Panther (Thomas, et al., 2003) using two unranked lists of genes (target and background lists).

### ATAC-seq

ATAC-seq was performed starting from 5000 sorted cells using a FAST-ATAC protocol previously described (Corces, et al., 2016). All the libraries were sequenced 2×50 bp with an Illumina Hiseq 2500 platform. We obtained at least 100 M reads from the sequencing of each library. For data comparison, we used two biological replicates for each condition. Reads were aligned to the danRer10 zebrafish genome and differential chromatin accessibility was calculated as reported (Magri, et al., 2019). All chromatin regions reporting a differential accessibility with an adjusted p-value < 0.05 were considered as differentially open chromatin regions (DOCRs). All the DOCRS have been associated with genes using the online tool GREAT (McLean, et al., 2010) with the option “basal plus extension”. Gene ontology analysis of all the genes associated with DOCRs was also performed with GREAT. De novo motif enrichment of TF binding sequences in the sets of DOCRs was performed using HOMER (Heinz, et al., 2010). Top enriched TF PWMs from the HOMER results and PWMs from JASPAR database (Fornes, et al., 2020) were used as input for the online tool FIMO (Grant, et al., 2011) to assess the exact TFBS genome position in the DOCRs. Before estimating the rate of TF co-occupancy in same peak among the binding motifs for the different TFs, all the binding motif sequences overlapping for more than 3 bps were eliminated, keeping only the TF binding sequences with the lowest p-value.

### Activating/repressing cis-regulatory element configuration

For this analysis only the DEGs and the DORCs that could be associated to each other were taken into account. The log2FoldChange values of transcript expression and chromatin accessibility of NR and RPE at 23 hpf were used to discriminate four clusters of activating and repressing CREs. Gene Ontology enrichment analysis of the genes belonging to the different clusters was performed with FishEnrichr (Kuleshov, et al., 2016).

### CRE 5mC analysis

Whole genome bisulfite sequencing and TET-assisted bisulfite sequencing (Bogdanovic, et al., 2016) data was trimmed with Trimmomatic software (Bolger, et al., 2014) and mapped onto the danRer10 genome assembly using WALT (Chen, et al., 2016) keeping only the reads mapping to a unique genomic location. Duplicates were removed with sambamba (Tarasov, et al., 2015) and DNA methylation levels were calculated using MethylDackel (https://github.com/dpryan79/MethylDackel). All heatmaps were made using Deeptools (Ramirez, et al., 2014) and Gene ontology enrichments were calculated with GREAT (McLean, et al., 2010). ATAC-seq reads were counted using BEDtools (Quinlan and Hall, 2010) and statistics were performed in R.

### CRISPR/Cas9 *F0 screening*

All the sgRNAs were designed using the online tool CRISPscan (https://www.crisprscan.org/) (Moreno-Mateos, et al., 2015) and synthetized following described protocols (Vejnar, et al., 2016). All the sgRNAs were selected to target the first half of the CDS in exons resulting actually expressed in the eye tissues from our RNA-seq data (trying to avoid the first exon to prevent the usage of an alternative start codon that would produce a possibly functional protein), with an efficiency score > 58 and no predicted off-targets (Table S3). Two different sgRNAs were used together to target the same gene. The sgRNAs were injected in the zebrafish yolk at 1-cell stage at final concentration of 80 ng/μl together with the Cas9 endonuclease at a concentration of 300 ng/μl. 1 nl of the mixture was injected in each embryo. In the case a target gene had a close paralogue, the sgRNAs targeting both of the paralogues were injected at the same time, adjusting the final concentration of the sgRNA-Cas9 mixture. Lethality, phenotypic features and penetrance were assessed at 24 and 48 hpf.

### Cell culture

Human hiPSCs were obtained, after informed consent, from peripheral blood monocytes by cell reprogramming using a non-integrative Sendai virus vector as described (Garcia Delgado, et al., 2019). Cells were maintained in feeder-free adherent conditions onto Matrigel-covered plates in standard incubation at 37°C, 5% CO_2_, 20% O_2_. hiPSCs were fed every two days with mTser1 serum-free culture medium and passaged every 5-7 days depending on confluency. Dispase was used for gentle dissociation for passage. A 6-well plate with undifferentiated, well-grown hiPSCs was the starting point of the experiment (Day 0). To induce RPE differentiation culture medium was changed to the following: KO DMEM, KSR 15%, Glutamax 2 mM, non-essential aminoacids 0.1 mM, β-mercaptoethanol 0.23 mM, Peniciline/streptomycin. Differentiating cells were harvested directly in Trizol LS (Invitrogen) at day 0, and weeks 1, 2, 3 and 4 for gene expression studies.

## Acknowledgements

We thank Lázaro Centanin, Juan Tena, Miguel Moreno-Mateos, Joaquin Letelier, Marta Magri, and Santiago Negueruela for their excellent scientific advice and critical input. Corin Díaz and Katherina García for their technical assistance with FACS. This work is supported by the following grants: (I) To JRMM: From the Spanish Ministry of Science, Innovation and Universities (MICINN): BFU2017-86339P with FEDER funds, and MDM-2016-0687. (II) To OB Australian Research Council (ARC) Discovery Project (DP190103852). (III) To FJDC: Andalusian Ministry of Health, Equality and Social Policies (PI-0099-2018). (IV) To PB: BFU2016-75412-R with FEDER funds; PCIN-2015-176-C02-01/ERA-Net Neuron ImproVision, and a CBMSO Institutional grant from the Fundación Ramón Areces. (V) To both JRMM and PB: BFU2016-81887-REDT, as well as Fundación Ramón Areces-2016 (Supporting LB).

## Author contribution

LB conducted most experiments, performed bioinformatic analyses, and had a main contribution in figures and manuscript edition. SN, TMM, and RP contributed to protocols refining, FACS experiments, and fish maintenance. BdC and FJDC carried out hiPSCs experiments together with LB. OB performed 5mC analysis and contributed to manuscript edition. JRMM conceived the project and assisted LB in data analysis. The manuscript was edited and written by JRMM and PB, who co-supervised all the work.

## Conflict of Interest

The authors declare that they have no conflict of interest

## Availability of Data

Datasets supporting the conclusions of this article are available in the Gene Expression Omnibus (GEO) repository (https://www.ncbi.nlm.nih.gov/geo) under the following accession numbers: RNA-seq (GSE150346) and ATAC-seq (GSE150189).

---

**Figure S1:**
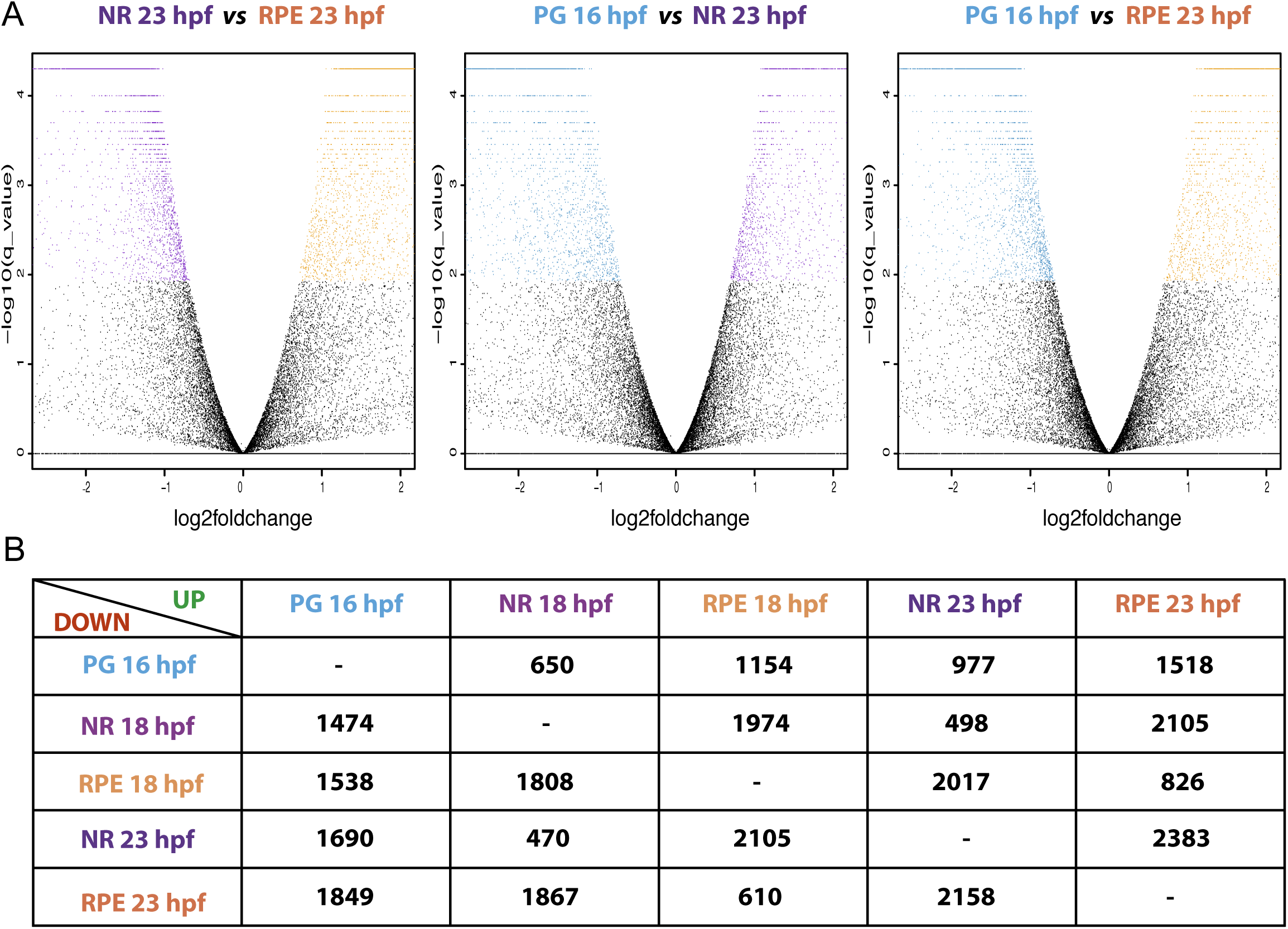
Eye domain transcriptome variations during optic cup morphogenesis. (A) Volcano plots illustrating the transcriptome variations during eye morphogenesis. Each dot corresponds to a gene. Black dots indicate not significant variations, whereas coloured dots point out significant expression variations among domains and developmental stages. (B) Table summarizing the number of DEGs (upregulated or downregulated) in each one of the conditions.

**Figure S2:**
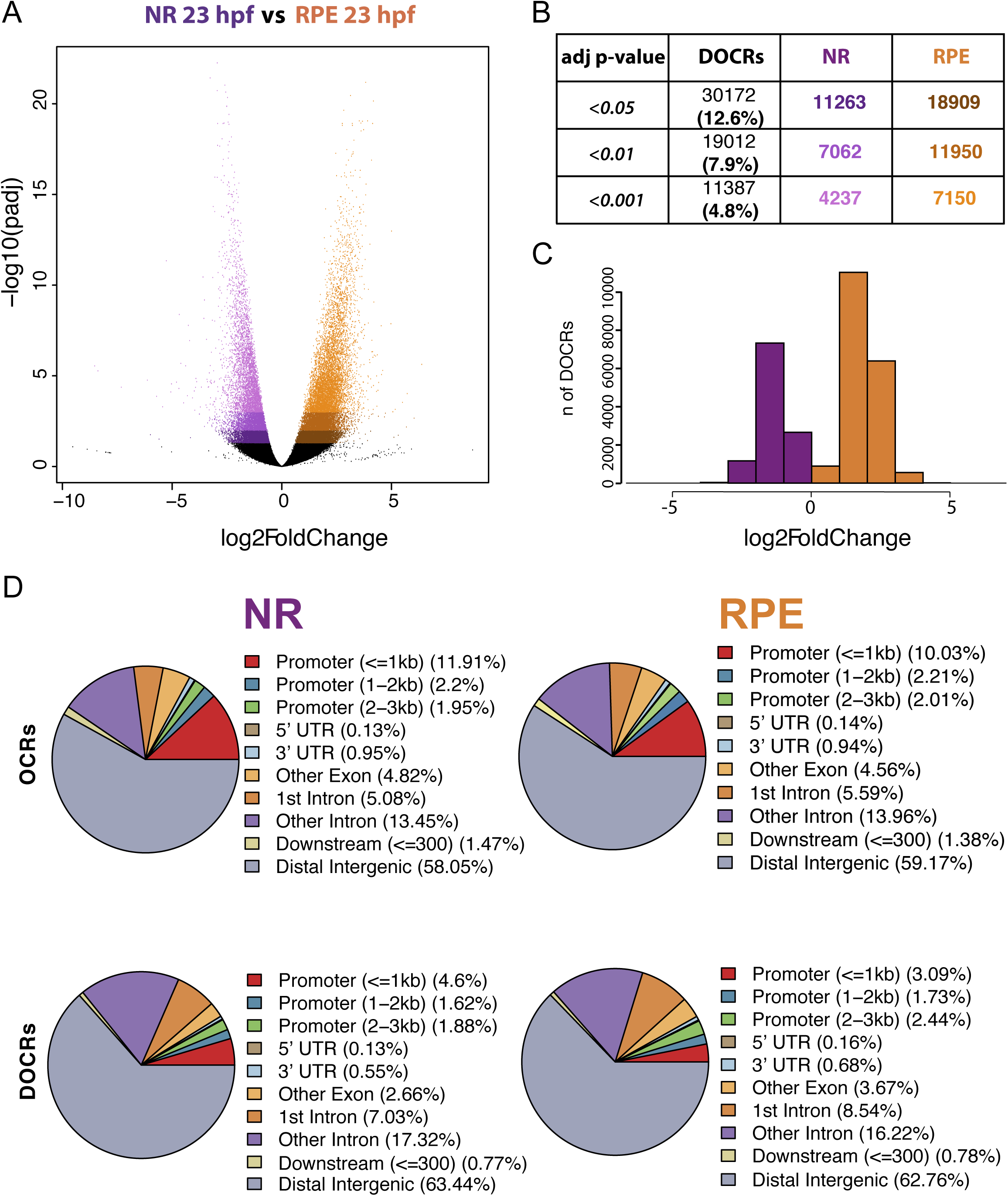
Analysis of differentially opened chromatin regions (DOCRs) during optic cup morphogenesis. (A) Volcano plot illustrating the chromatin accessibility changes during eye morphogenesis. Each dot corresponds to a peak (i.e. open chromatin region). Black dots indicate not significant variations; colour shades chromatin accessibility changes with different ranges of adjusted p-value (darker: p<0.05; medium= p<0.01, lighter= p<0.001). (B) Table including the number of peaks significantly more or less accessible between the two conditions. (C) Frequency histogram showing the distribution of the DOCRs in relation to their accessibility fold change. (D) Pie charts displaying the percentage of peaks falling in distinct regions of zebrafish genome. Top: genome distribution of the whole set of open chromatin regions (OCRs) identified by ATAC-seq in each condition. Bottom: genome distribution of the DOCRs.

**Figure S3:**
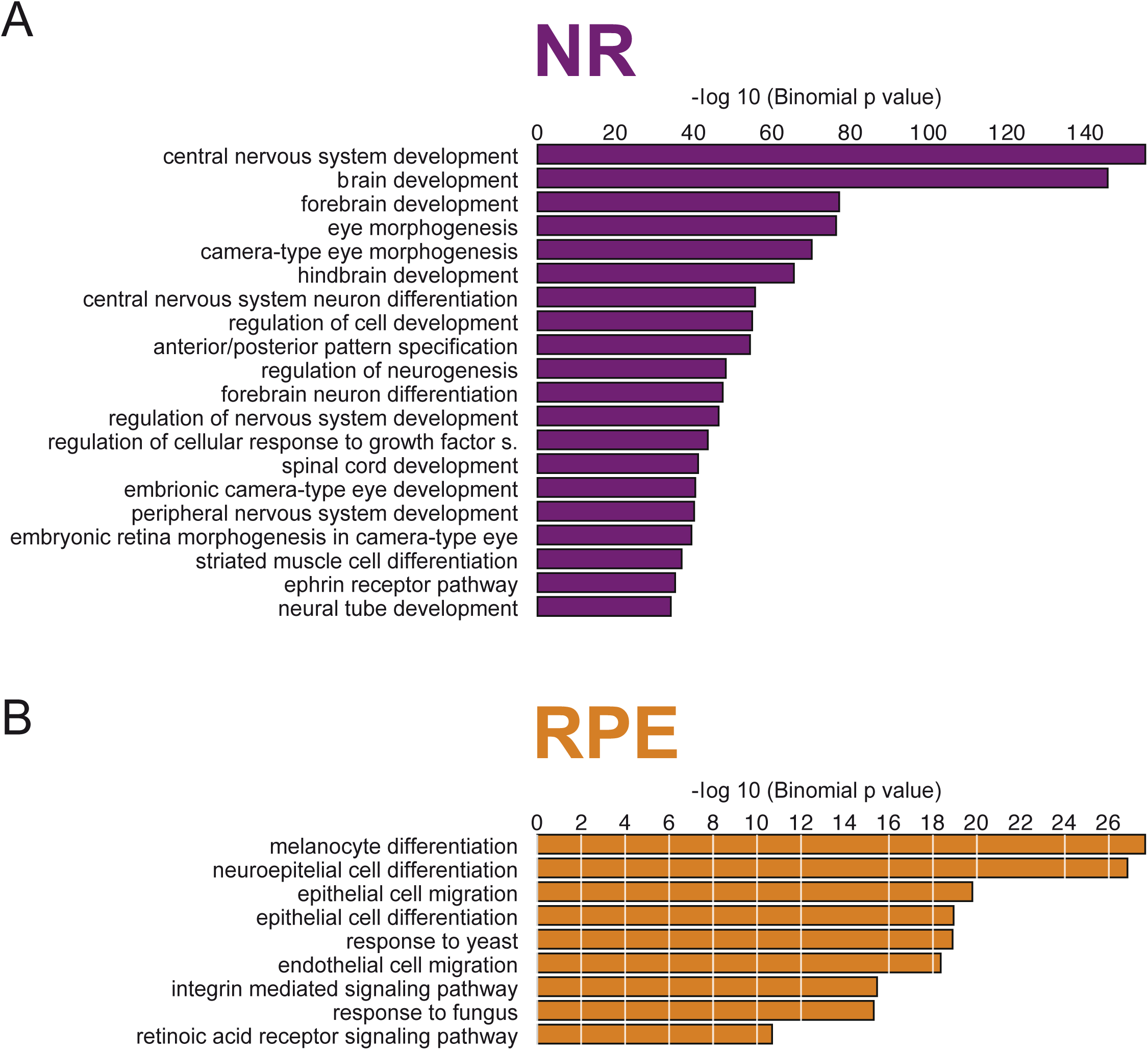
Gene ontology enrichment of the genes associated with DOCRs. Bar chart showing GO terms for biological processes enriched in the genes associated to DOCRs. Bar length is proportional to enrichment significance.

**Figure S4.**
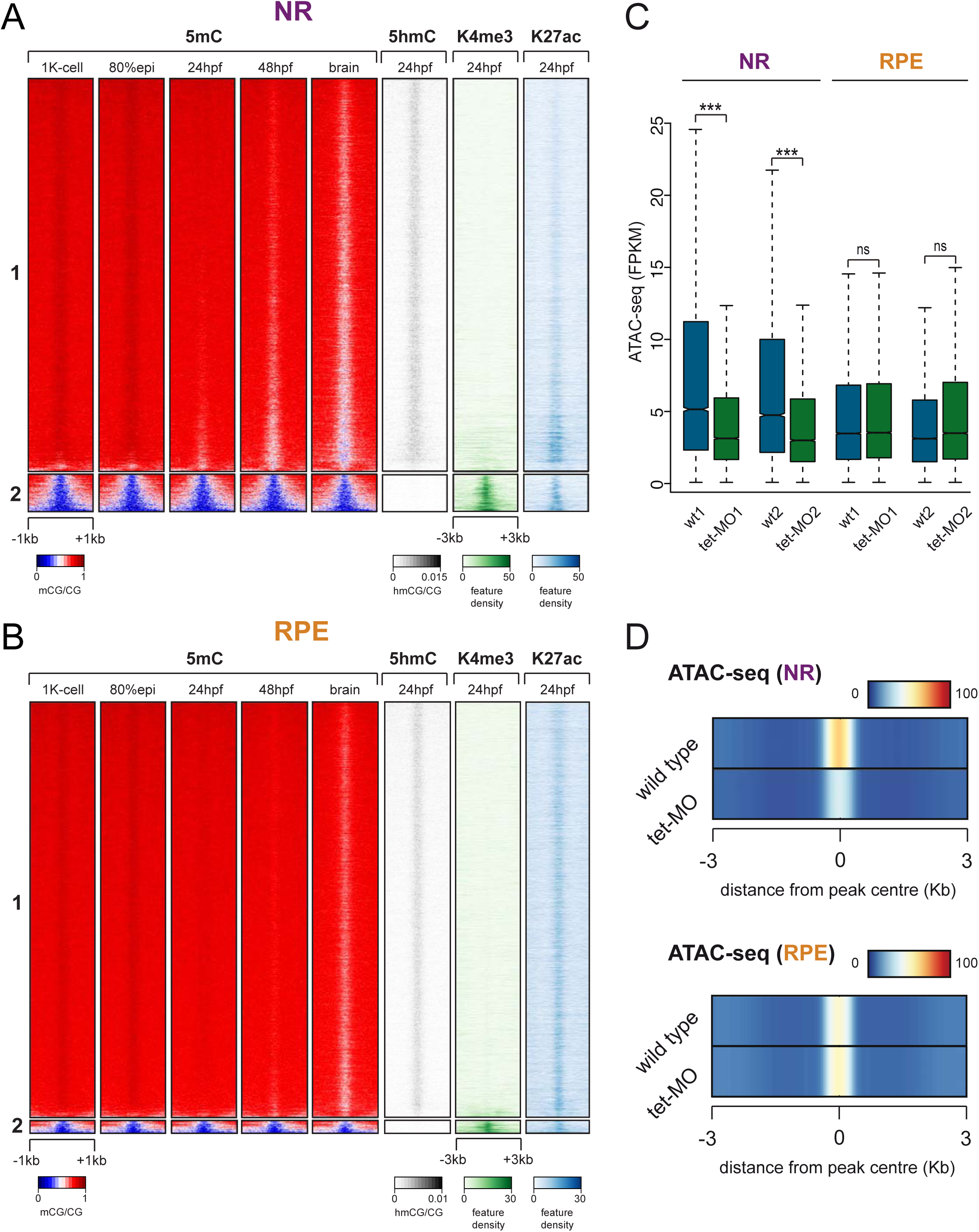
DNA methylation profiles of NR and RPE peaks. Clustering of NR (A) and RPE (B) ATAC-seq peaks according to 5mC pattern across developmental stages and in adult brain (left columns), as well as 5hmC, K4me3, and K27ac profiles at 24 hpf (right columns). Note that the signature of cluster #1 corresponds to active enhancers, whereas that of cluster #2 corresponds to hypomethylated promoter regions. (C) Chromatin accessibility quantification for NR and RPE peaks in wild type and triple tet (tet1/2/3) morphants. Analysis was performed in whole 24hpf embryos. Statistical significance was evaluated using a Mann-Whitney-Wilcoxon Test, P < 2.2e-16. (D) Chromatin accessibility levels for NR and RPE ATAC-seq peaks in wild type and and triple tet (tet1/2/3) zebrafish morphants.

**Figure S5:**
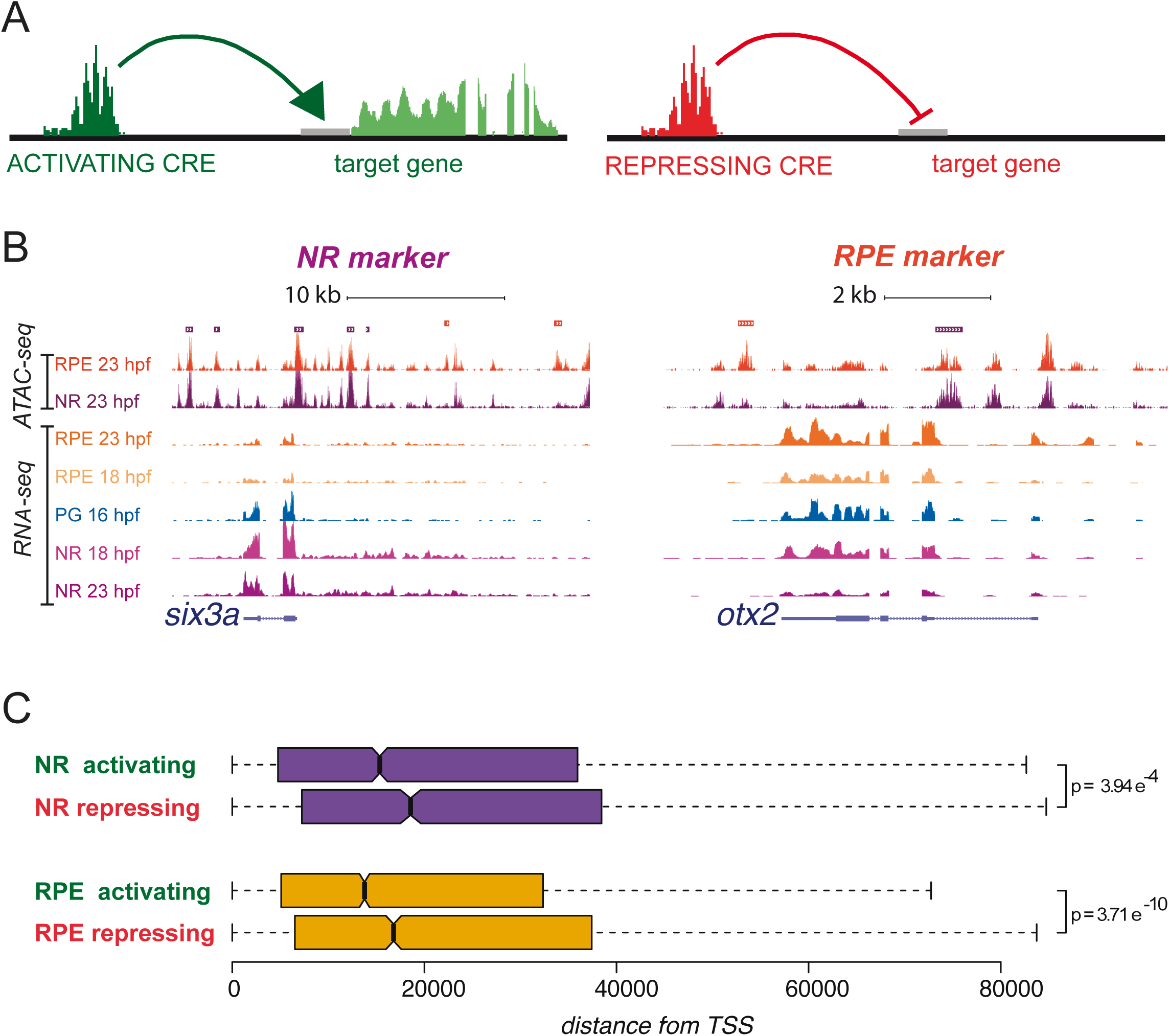
Activating and repressing CREs localization. (A) Schematic representation of the functional relations among DOCRs, either activating (green) or repressing (red), and their associated DEGs. (B) Illustrative examples of ATAC-seq and RNA-seq tracks (UCSC browser) for representative NR (six3; E) and RPE (otx2; F) genes. Bars on the top indicate DOCRs. If purple, the DOCR is more accessible in NR; if orange, the DOCR is more accessible in RPE. Some of the more accessible DOCRs are accompanied by a decreased transcription of the associated gene in the corresponding tissue. (C) Average distance to the TSS of all activating and repressing DOCRs for the NR and RPE domains (see Figure 2C). Note that activating CREs are significantly (T-test) closer to the TSS than repressing regions in both domains.

**Figure S6:**
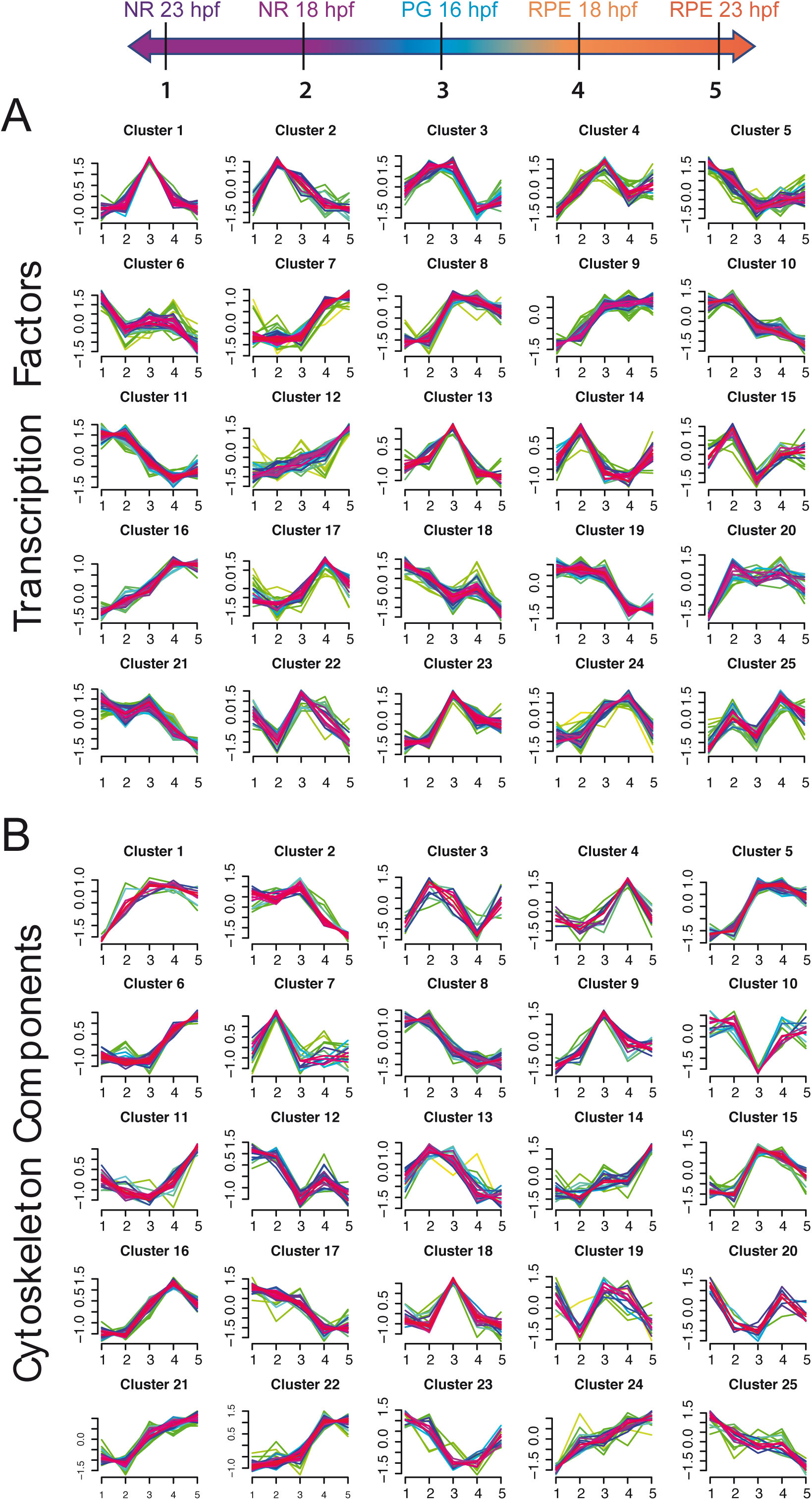
Partitioning clustering of gene expression variations during optic cup development. Partitioning clustering output (k= 25) showing the expression trends in the distinct domains and stages examined for two classes of genes: TFs (A) and cytoskeleton components (B). Stages and domains examined are linearly represented with a colour code and numbered from 1 (NR 23hpf) to 5 (RPE 23hpf), following the lineage trajectory.

**Figure S7:**
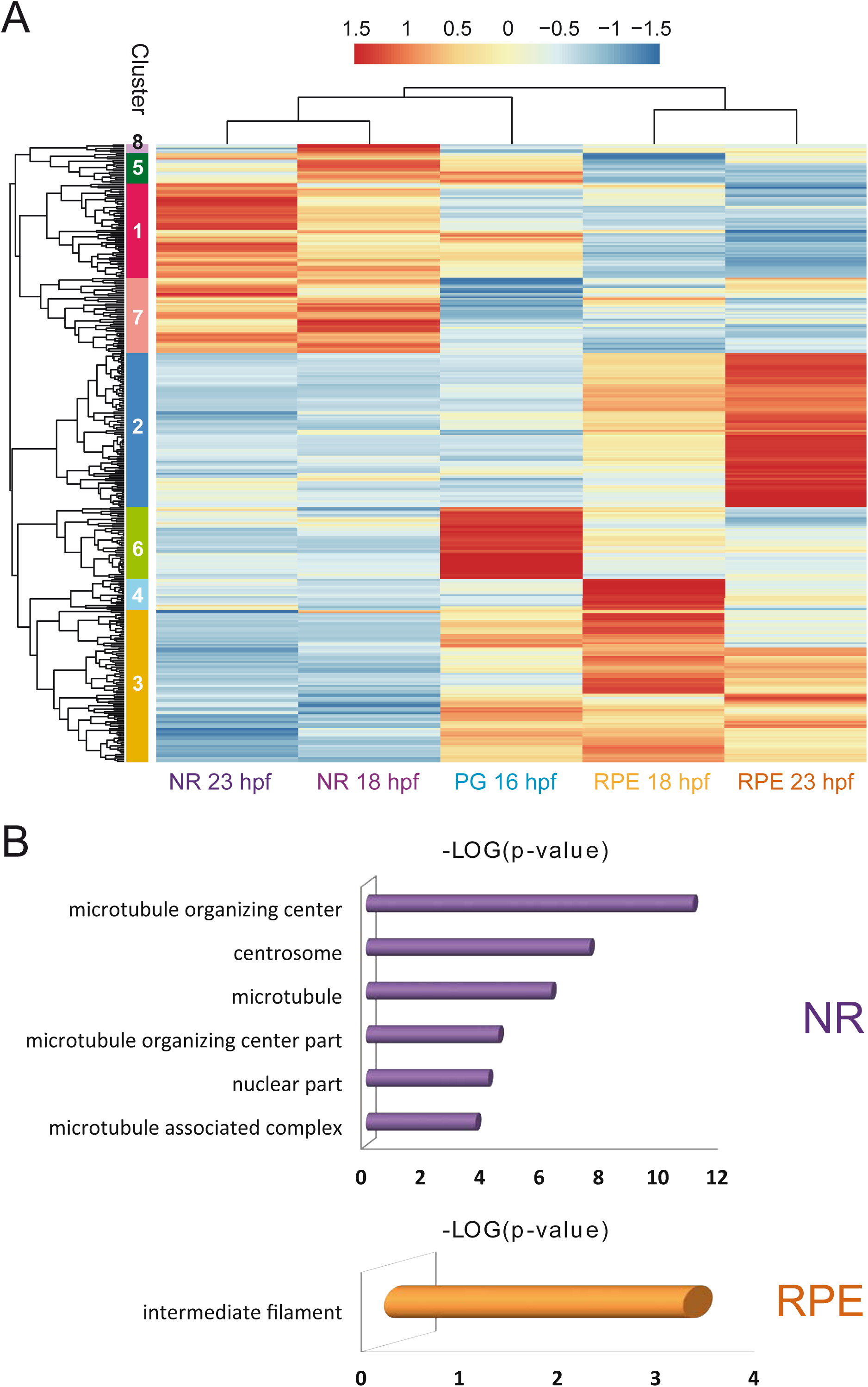
Hierarchical clustering of gene expression variations during optic cup development for cytoskeletal components. (A) Hierarchical clustering output showing the expression trends in distinct domains and stages. Gene expression values, normalized by row, are indicated with a red to blue graded colour. Note that all stages and domains are represented by at least one gene cluster (B) Cellular component GO enrichment for cytoskeletal genes belonging to NR or RPE clusters.

**Figure S8:**
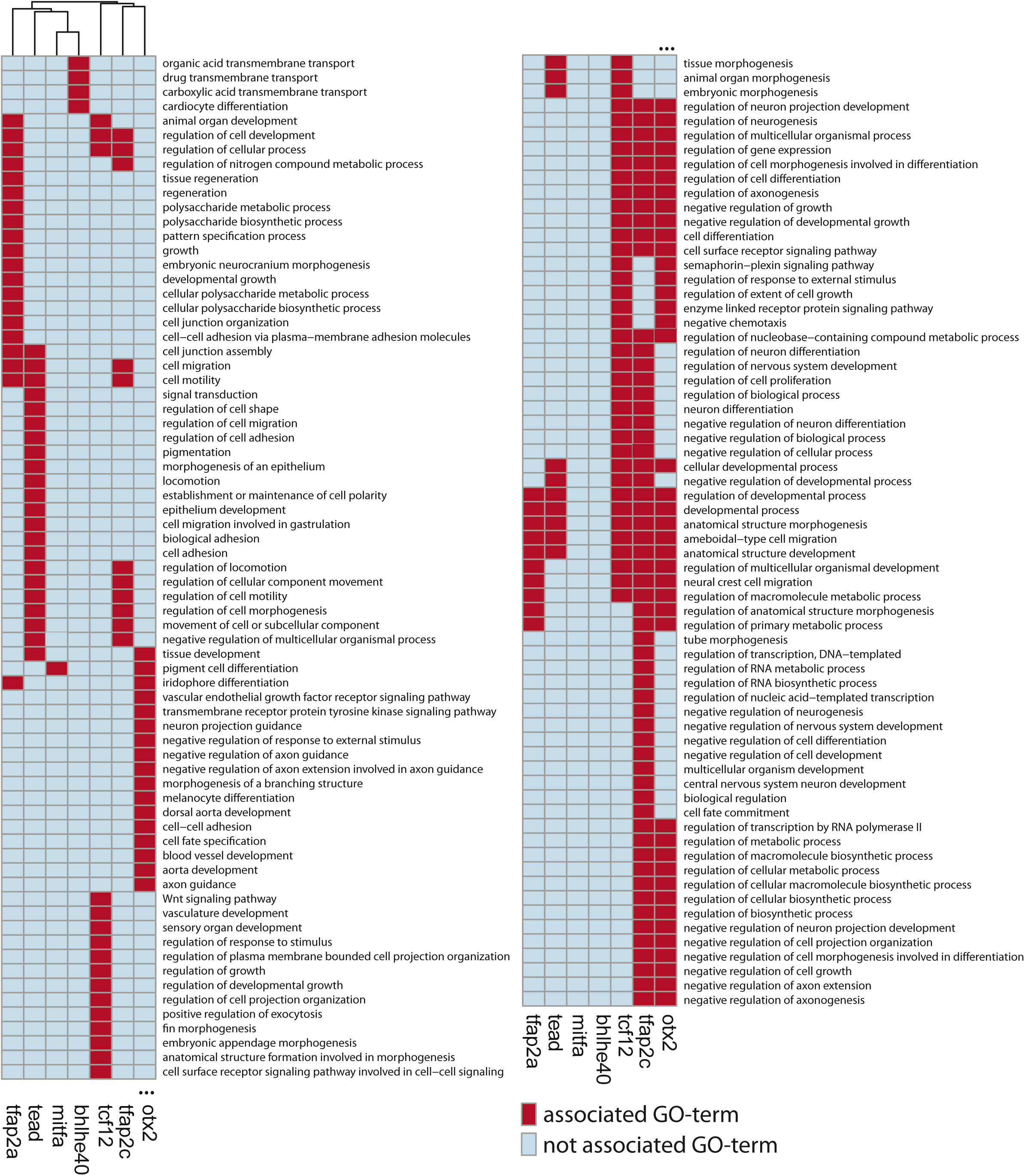
GO terms associated with RPE TFs. Hierarchical clustering of the main RPE TFs according to the enriched GO terms (Biological process) associated to their putative downstream genes (genes associated with RPE DOCRs that contain each TFBS).

**Figure S9:**
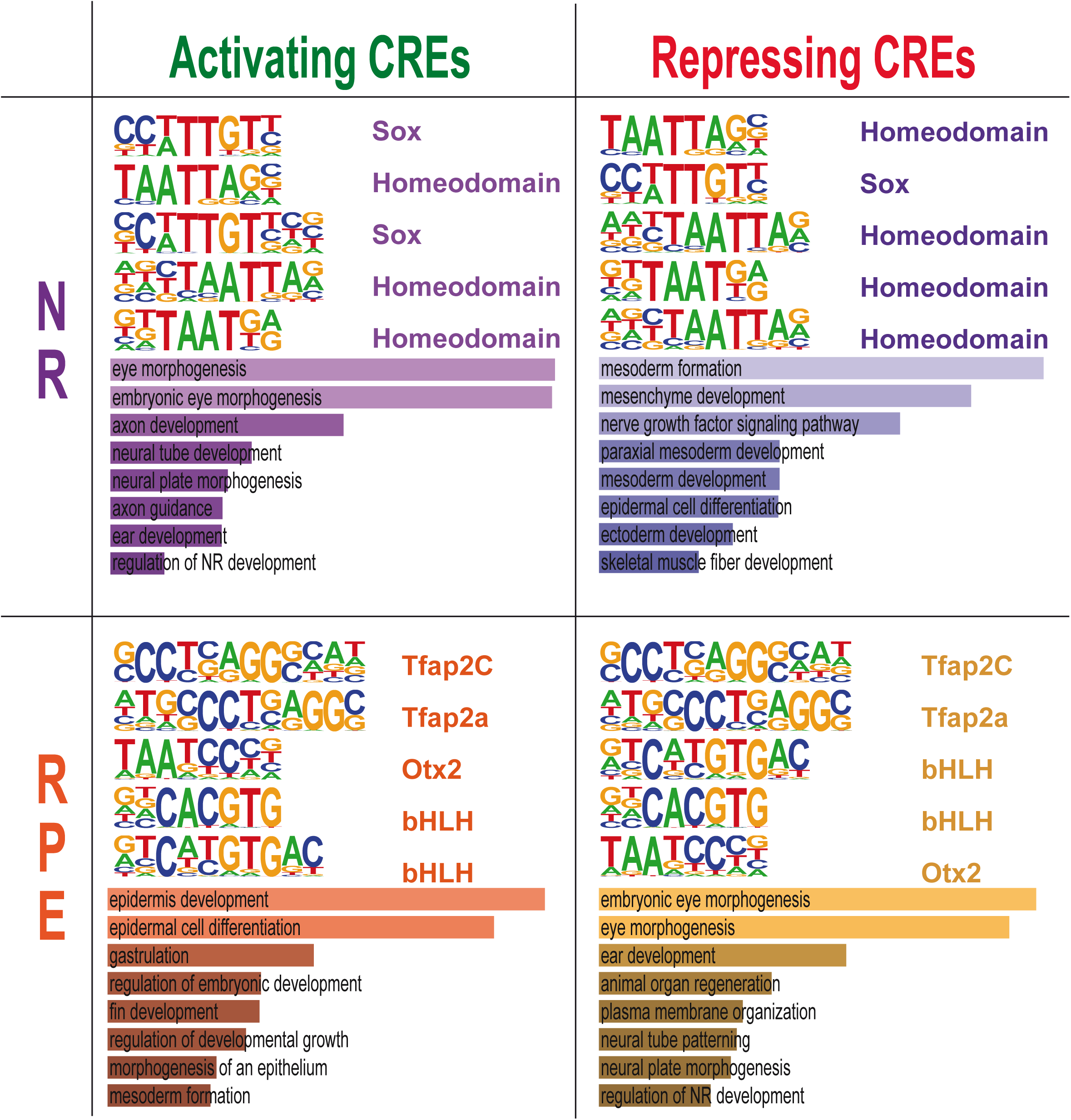
Motif enrichment analysis of activating and repressing CREs. Representative TF motifs enriched in activating (left column) and repressing (right column) CREs in both the NR (A) and RPE (B) domains. Analysis of GO terms enrichment for genes associated with each set of elements is indicated.

**Figure S10:**
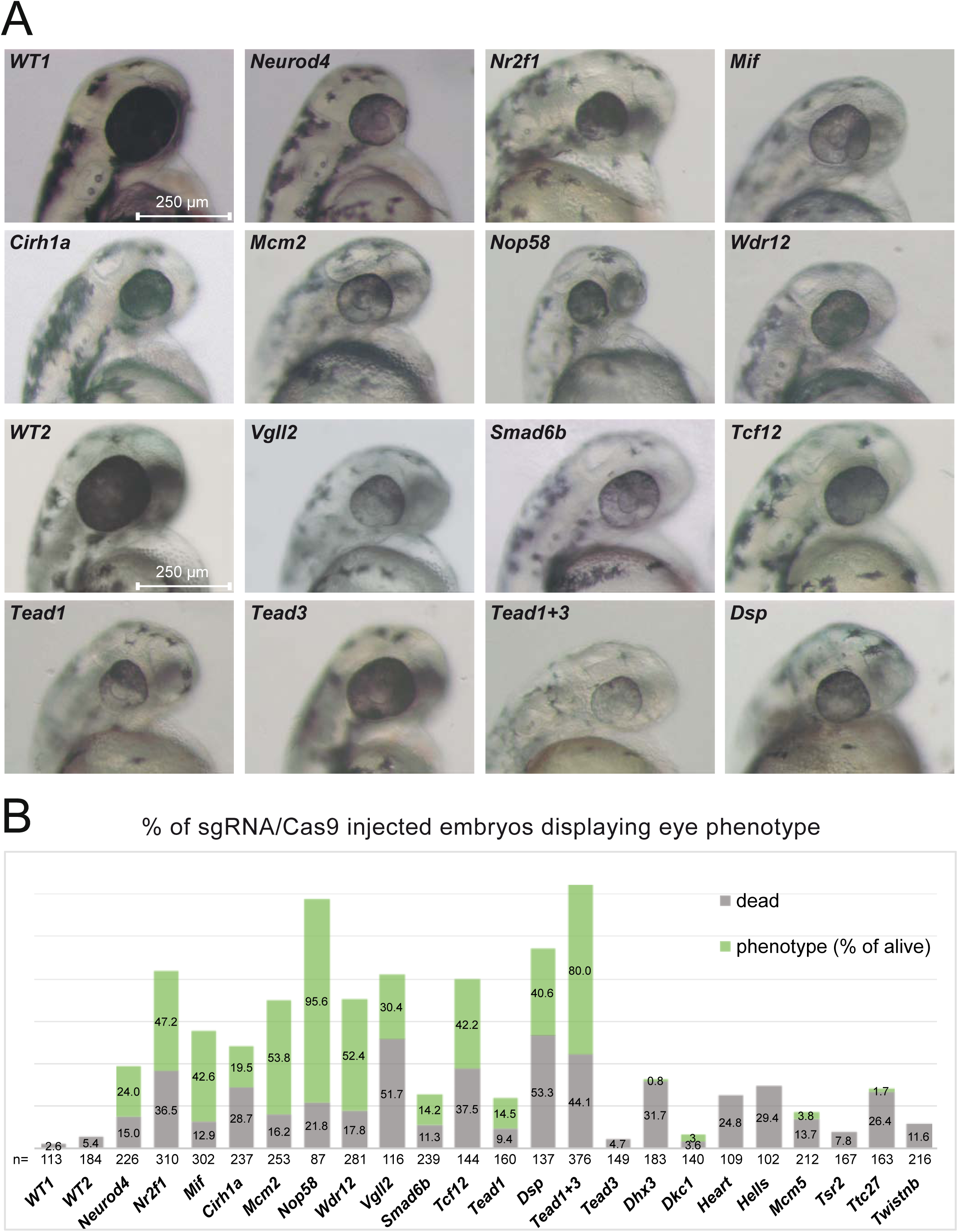
CRISPR CRISPR/Cas9 F0 screening. (A) Eye phenotypes resulting from the injection of Cas9 protein and sgRNAs for the candidate genes. (B) Percentage of embryos showing an impaired eye phenotype upon the injection of Cas9/sgRNAs complex (n= total number of injected embryos). The % of lethality upon injection is also shown. See also Supplementary data set 12.

## Appendix Supplementary Tables

**Table SI.**
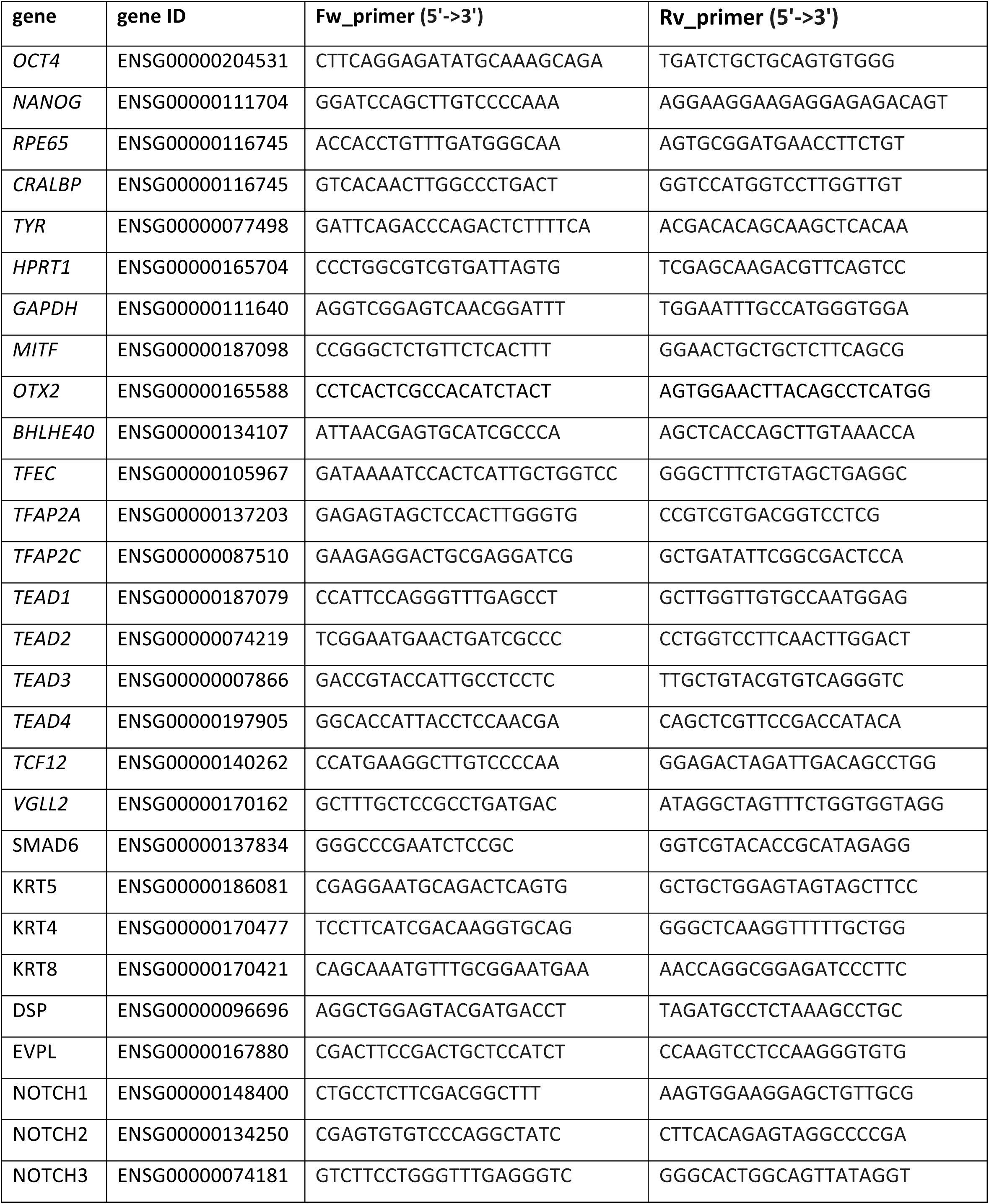
Primers used for Real Time q-PCR of human samples.

**Table SII.**
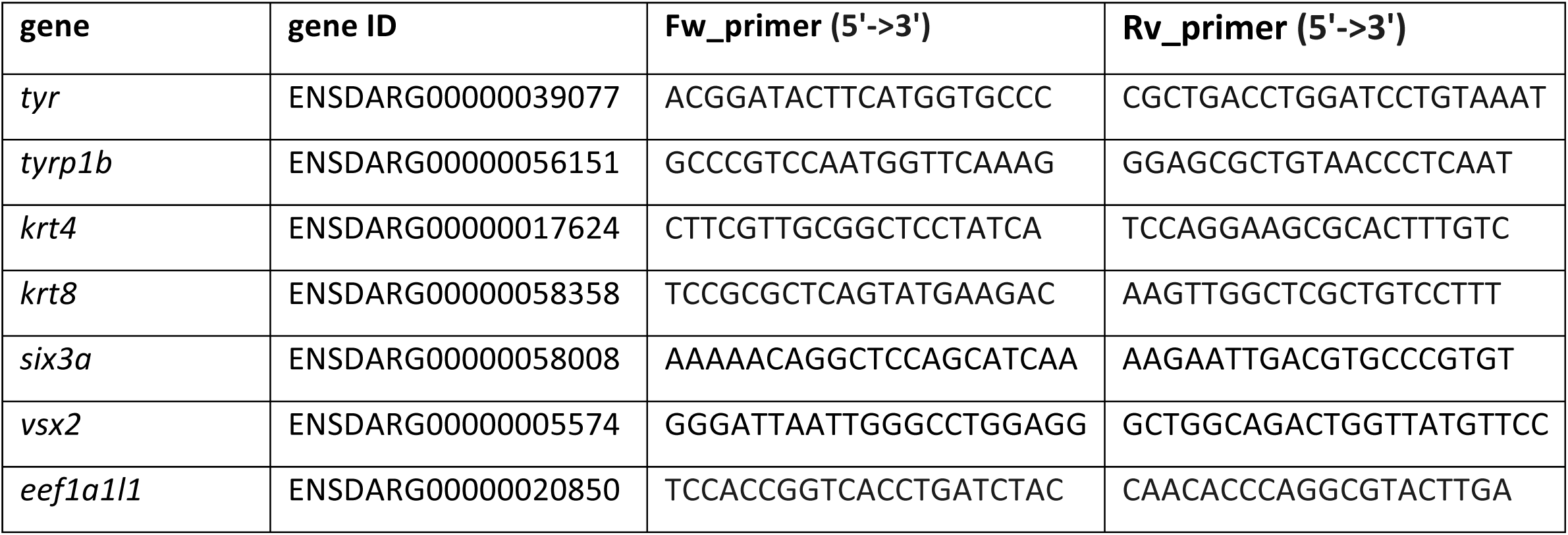
Primers used for Real Time q-PCR of zebrafish samples.

**Table SIII.**
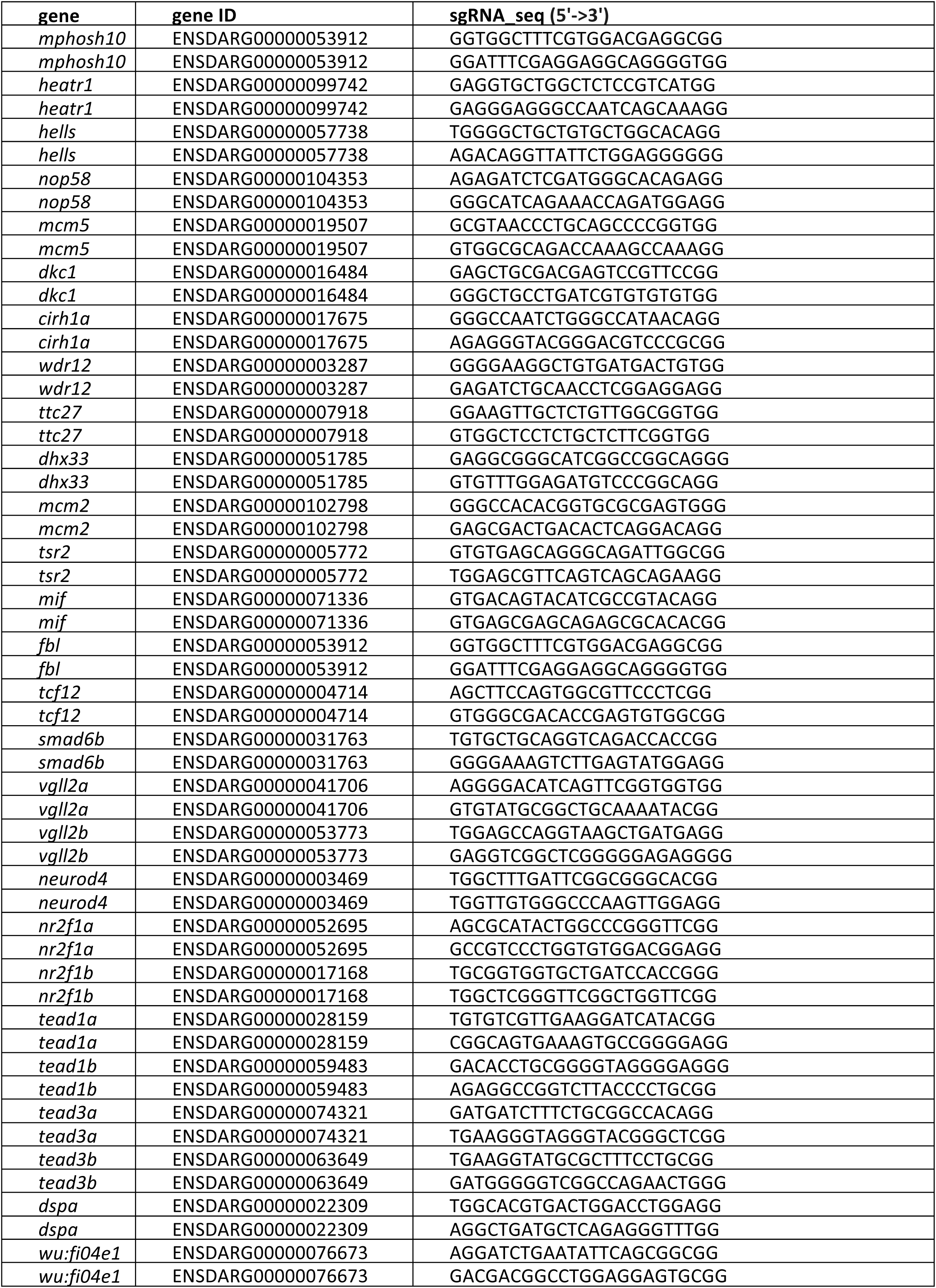
sgRNAs used for CRISPR/Cas9 screen

## Notes

### Competing Interest Statement

The authors have declared no competing interest.

